# The fatty acid site is coupled to functional motifs in the SARS-CoV-2 spike protein and modulates spike allosteric behaviour

**DOI:** 10.1101/2021.06.07.447341

**Authors:** A. Sofia F. Oliveira, Deborah K. Shoemark, Amaurys Avila Ibarra, Andrew D. Davidson, Imre Berger, Christiane Schaffitzel, Adrian J. Mulholland

## Abstract

The SARS-CoV-2 spike protein is the first contact point between the SARS-CoV-2 virus and host cells and mediates membrane fusion. Recently, a fatty acid binding site was identified in the spike (Toelzer *et al*. Science 2020). The presence of linoleic acid at this site modulates binding of the spike to the human ACE2 receptor, stabilizing a locked conformation of the protein. Here, dynamical-nonequilibrium molecular dynamics simulations reveal that this fatty acid site is coupled to functionally relevant regions of the spike, some of them far from the fatty acid binding pocket. Removal of a ligand from the fatty acid binding site significantly affects the dynamics of distant, functionally important regions of the spike, including the receptor-binding motif, furin cleavage site and fusion-peptide-adjacent regions. The results also show significant differences in behaviour between clinical variants of the spike: e.g. the D614G mutation shows a significantly different conformational response for some structural motifs relevant for binding and fusion. The simulations identify structural networks through which changes at the fatty acid binding site are transmitted within the protein. These communication networks significantly involve positions that are prone to mutation, indicating that observed genetic variation in the spike may alter its response to linoleate binding and associated allosteric communication.

## Introduction

The COVID-19 pandemic, which is having a devastating social and economic impact worldwide, is caused by the severe acute respiratory syndrome 2 (SARS-CoV-2) coronavirus. Since the initial outbreak in late 2019, SARS-CoV-2 has caused >156 million confirmed cases of COVID-19 disease and >3.2 million deaths (*1*) (perhaps as many as 7-13 million (*2*)) worldwide as of 8^th^ May, 2021. SARS-CoV-2 is an enveloped, single-stranded RNA virus that belongs to the *Betacoronavirus* genus of the *Coronaviridae* family which includes pathogenic human coronaviruses that cause SARS (severe acute respiratory syndrome) and MERS (Middle East respiratory syndrome) (*3, 4*). It initially infects respiratory epithelial cells by binding to the angiotensin-converting 2 enzyme (ACE2) (*5, 6*). When it first emerged as a human pathogen, SARS-CoV-2 was thought to cause predominantly respiratory disease, particularly pneumonia and severe acute respiratory distress syndrome (*7, 8*). However, it is now known that its effects are not limited to the respiratory tract: COVID-19 can cause severe inflammation and damage in other organs (*9-11*), including the heart, kidneys, liver and intestines, and can lead to neurological problems (*12*). In common with other enveloped viruses, SARS-CoV-2 fuses its viral envelope with a host cell membrane to infect cells. Membrane attachment and fusion with the host cell is mediated by the SARS-CoV-2 spike protein, which primarily binds to the host ACE2 receptor (*5, 6*); however, the spike can also interact with neuropilin-1 (*13*) and potentially with other receptors (*14*). The spike is a glycoprotein (*15, 16*) and is found on the surface of the virion, in a trimeric form (Figure 1A).

**Figure 1.**
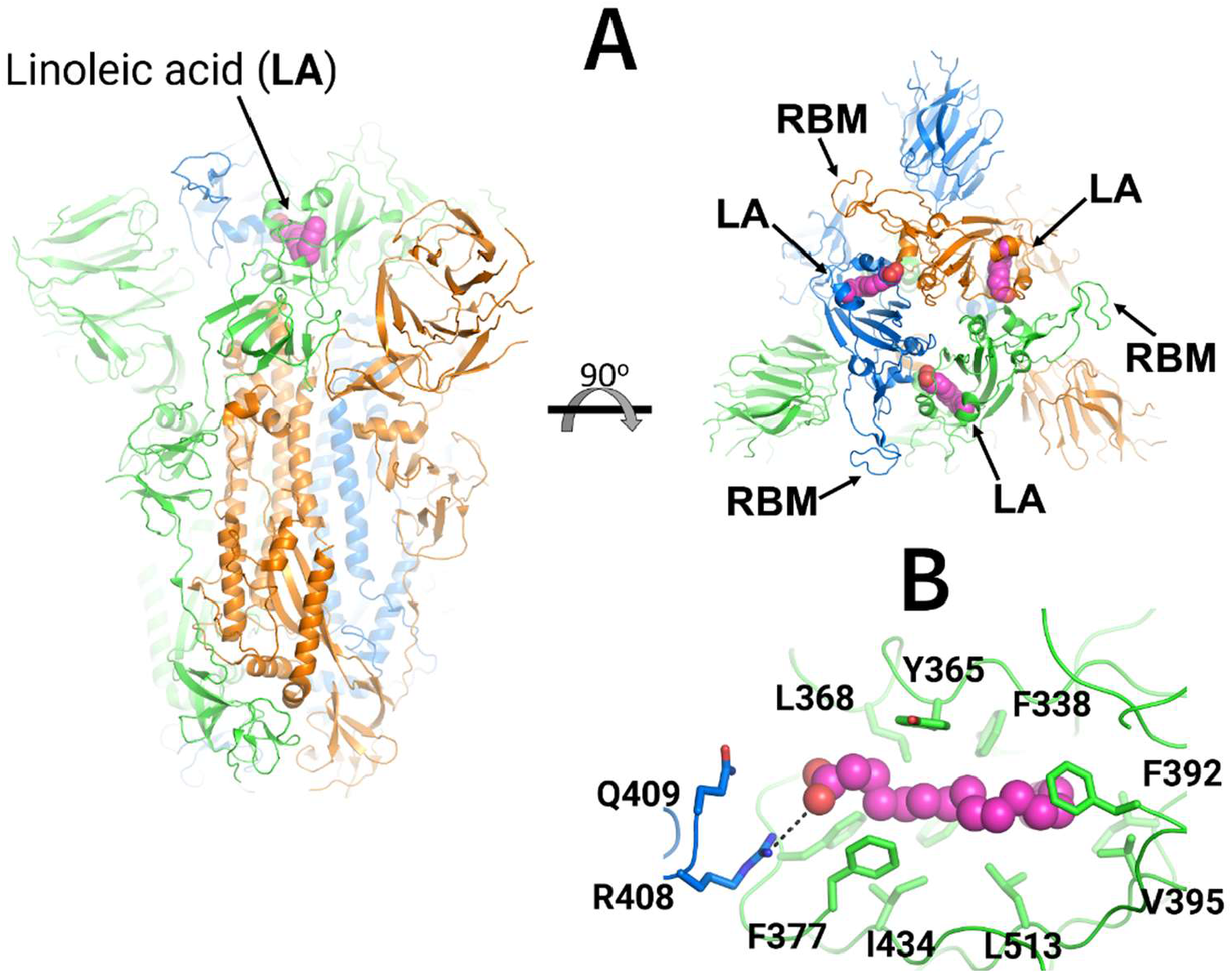
Cryo-EM structure of the ectodomain of the SARS-CoV-2 spike trimer with linoleic acid (LA) bound to the fatty acid-binding sites (*17*). (**A**) Three-dimensional structure of the complex of the locked (all RBMs occluded) ectodomain of the SARS-CoV-2 spike trimer with linoleic acid (PDB code: 6ZB5) (*17*). The spike protein is a homotrimer (*16*): each monomer is shown in a different colour, namely green, orange and blue. LA molecules are highlighted with spheres. Each fatty acid (FA) binding site is located at the interface between two neighbouring monomers, and is formed by residues from two adjacent receptor-binding domains. (**B**) Detailed view of the FA binding site: this pocket is lined by hydrophobic and aromatic residues and the LA acidic headgroup is close to R408 and Q409.

Each monomer is formed of three domains: a large ectodomain, a transmembrane anchor and a short cytoplasmic tail (*16*). The ectodomain comprises two subunits: S1 is responsible for binding to ACE2 (*6, 16*), and S2 for viral-host membrane fusion (*16, 18*). The SARS-CoV-2 spike contains two proteolytic cleavage sites (*16*): one ‘furin protease recognition’ site at the S1/S2 boundary, thought to activate the protein (*19*), and a second in the S2 subunit (S2’
s) that releases the fusion peptide (*16, 18*). The SARS-CoV-2 spike contains three free fatty acid (FA) binding sites, each located at the interface between every two neighbouring receptor-binding domains (Figures 1A and S3) (*17*). The FA binding sites are lined by aromatic and hydrophobic residues (Figure 1B) and a positively charged residue from a neighbouring monomer, namely R408, which acts as an anchor for the FA carboxylate headgroup (*17*). The open spike conformation, with at least one RBD pointing upwards, is needed to interact with ACE2 receptors on the human host cell. It was shown by surface plasmon resonance that the presence of the FA linoleic acid (LA) reduces binding of the spike to ACE2 (*17*). In agreement, LA stabilises the locked spike conformation, in which the RBM is occluded and cannot bind to the human ACE2 receptor (*17*), but there is no obvious connection between the FA sites and other structural motifs relevant for membrane fusion, or with antigenic epitopes. MD simulations showed persistent and stable interactions between LA and the spike trimer (*17, 20*). These simulations also revealed that LA rigidifies the FA binding site, and these effects extend to the N-terminus domain (*20*). The cryo-EM structure of the spike from pangolin coronavirus (which is closely related to SARS-CoV-2) shows that the spike also binds LA in an equivalent FA pocket (*21*). An equivalent FA binding site was also found on the Novavax SARS-CoV-2 construct expressed and purified from insect cells (*22*).

Simulations, particularly atomistic molecular dynamics (MD) simulations, have provided crucial atomic-level insight into the structure, dynamics and interactions of the SARS-CoV-2 spike (*13, 17, 20, 23-28*). Here, we apply dynamical-nonequilibrium MD simulations (*29-31*) to investigate the response of the SARS-CoV-2 spike to LA removal. We have shown this approach to be effective in identifying structural communication pathways in a variety of proteins, e.g. in identifying a general mechanism of interdomain signal propagation in nicotinic acetylcholine receptors (*32, 33*) and mapping the networks connecting the allosteric and catalytic sites in two clinically relevant β-lactamase enzymes (*34*). This approach is based on, first of all, equilibrium simulations of the system in question, which generate configurations for multiple dynamical-nonequilibrium simulations where the effect of a perturbation can be studied. Running a large number of nonequilibrium simulations allows for the determination of the statistical significance of the structural response observed.

### Dynamic response of the wild-type spike

A model of the locked wild-type spike was created from the cryo-EM structure (PDB code: 6ZB5) of the SARS-CoV-2 spike protein bound to three linoleate molecules (*17*). Missing loops were built to generate the wild-type sequence according to the Uniprot accession number P0DTC2 for the unglycosylated ectodomain of the spike bound with LA (for details, see Supplementary Material). The locked structure had 42 disulphides per trimer that remained intact and faithfully retained the structure and overall fold of the cryo-EM structure over the equilibrium simulation time (*17, 20*). Three equilibrium MD simulations (Figure S1), 200 ns each, were performed for the locked form of the unglycosylated and uncleaved (no cleavage at the S1/S2 interface) ectodomain of the spike bound with LA and used as starting points for 90 dynamical-nonequilibrium simulations (Figure S2). Here we have used models of the uncleaved spike ectodomains in order to detect any potential effects on structurally distant sites influenced by ligand in the FA sites in the intact spike. In these nonequilibrium simulations, all LA molecules were (instantaneously) annihilated. This triggers a response of the protein, as it adapts to LA removal (top panel in Figure S2). This annihilation is carried out for multiple configurations sampled from equilibrium MD, and the comparison between the equilibrium and short dynamical-nonequilibrium MD trajectories identifies the structural response of the protein. Running multiple (in this case, 90) dynamical-nonequilibrium simulations reduces the noise associated with the structural response of the protein and allows for the determination of the statistical significance of the observed response. Nonequilibrium simulations of this type are emerging as an effective tool to study signal transmission and identify communication networks within proteins (*32-36*). Here, the direct comparison between the equilibrium LA-bound and nonequilibrium apo spike simulations using the Kubo-Onsager approach (*29-31*) (bottom panel in Figure S2), and the average of the results over all the 90 replicates, allows for identification of the temporal sequence of conformational changes associated with the response of the spike to LA removal (Figures 2 and S4), and also the determination of their statistical significance (Figure S5). The structure that we simulate here corresponds to the unglycosylated wild-type spike (Uniprot accession number P0DTC2), not cleaved at the ‘furin protease recognition’ site. It was built based on the cryo-EM structure that originally revealed the FA binding site (PDB code: 6ZB5) (*17*). Although a few glycans (e.g. at positions N165, N234, and N343) have been shown to be involved in the spike infection mechanism by altering the dynamics of receptor binding domain opening (*23, 37*), and the glycan shield plays a vital role in the biological function of the spike, the internal networks and response of the protein scaffold identified here are not likely to be qualitatively altered by the glycans, which predominantly cover the exterior of the spike. As Casalino *et al*. have shown, glycan dynamics are fast relative to the dynamics of the protein (*23*). Note that the perturbation introduced here (LA annihilation) is not intended to mimic the physical process of LA (un)binding, but rather to promote a rapid response and force signal transmission within the protein, thus explicitly mapping the mechanical and dynamic coupling between the structural elements involved in this response. Note also that, due to the non-physical nature of the perturbation, the timescales observed for the protein’s response do not represent the physical timescales of conformational change, however, the responses of similar systems (e.g. wild-type and D614G spike) can be meaningfully compared.

**Figure 2.**
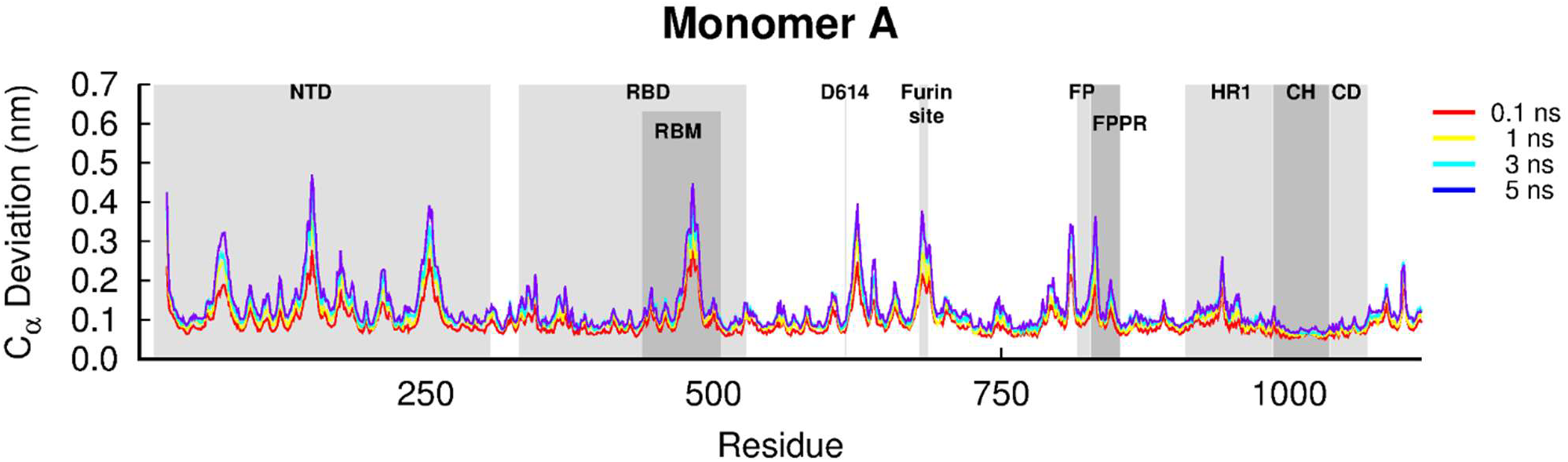
Average Cα-positional deviation for the first monomer in the five nanoseconds after LA removal from the FA sites in the SARS-CoV-2 spike. The structural deviations were calculated using the Kubo-Onsager approach (*29-31*) for the pairwise comparison between the equilibrium LA-bound and dynamical-nonequilibrium apo spike simulations and averaged over all 90 replicates. A similar response to LA removal is observed for the other two monomers (Figure S4). Some relevant motifs are highlighted in grey, namely N-terminal domain (NTD), receptor-binding domain (RBD), receptor-binding motif (RBM), fusion peptide (FP), fusion-peptide proximal region (FPPR), heptad repeat 1 (HR1), central helix (CH) and connector domain (CD).

Despite small variations in amplitude, the structural response to LA removal is similar for the three monomers (Figures 2 and S4). Figures 2 and S4 show the time evolution of the average Cα-positional deviation of each individual monomer in the 5 ns following LA. Comparing these figures shows that all the monomers respond similarly, with the same motifs and order of events associated with signal propagation observed for each.

The simulations show that the structural response to LA removal occurs in very specific and well-defined regions of the spike. It is striking that some functional motifs, including regions distant from the FA site, are particularly affected by LA removal (Figures 2 and S4). Upon LA removal, the response within the spike starts in the FA binding pocket region, and it rapidly propagates through the receptor-binding domain (RBD) to the N-terminal domain (NTD), furin site and residues surrounding the fusion peptide (FP). All of the monomers are closely intertwined, and therefore signal propagation does not occur simply within an individual monomer, but rather involves a complex network of conformational changes spanning all three chains (Figures S6-S8). For example, LA removal from FA pocket (formed by monomers A and C) induces structural responses in the NTD of monomer B (NTDB), RBD of monomer C (RBDC) and the furin site of monomer A, as shown by the propagation of the signal from the first site to these regions (Figures S6-S8).

### FA sites modulate key motifs for membrane fusion or antigenic epitopes in the spike

The RBMs respond rapidly to LA removal (Figures 3 and S9-S10), due to their close proximity to the FA sites. After 0.1 ns, significant structural rearrangements are already apparent in the RBMs, mainly in the A475-C488 segment (Figures 3 and S9-S10). Subsequently, a gradual increase in deviations is observed for A475-C488. The RBM lies between the β4 and β7 strands of the RBD and contains most of the residues that directly interact with ACE2 (*38, 39*). This motif is one of the most variable regions of SARS-CoV-2 spike (*40*) and a major target for neutralising antibodies (*41-43*).

**Figure 3.**
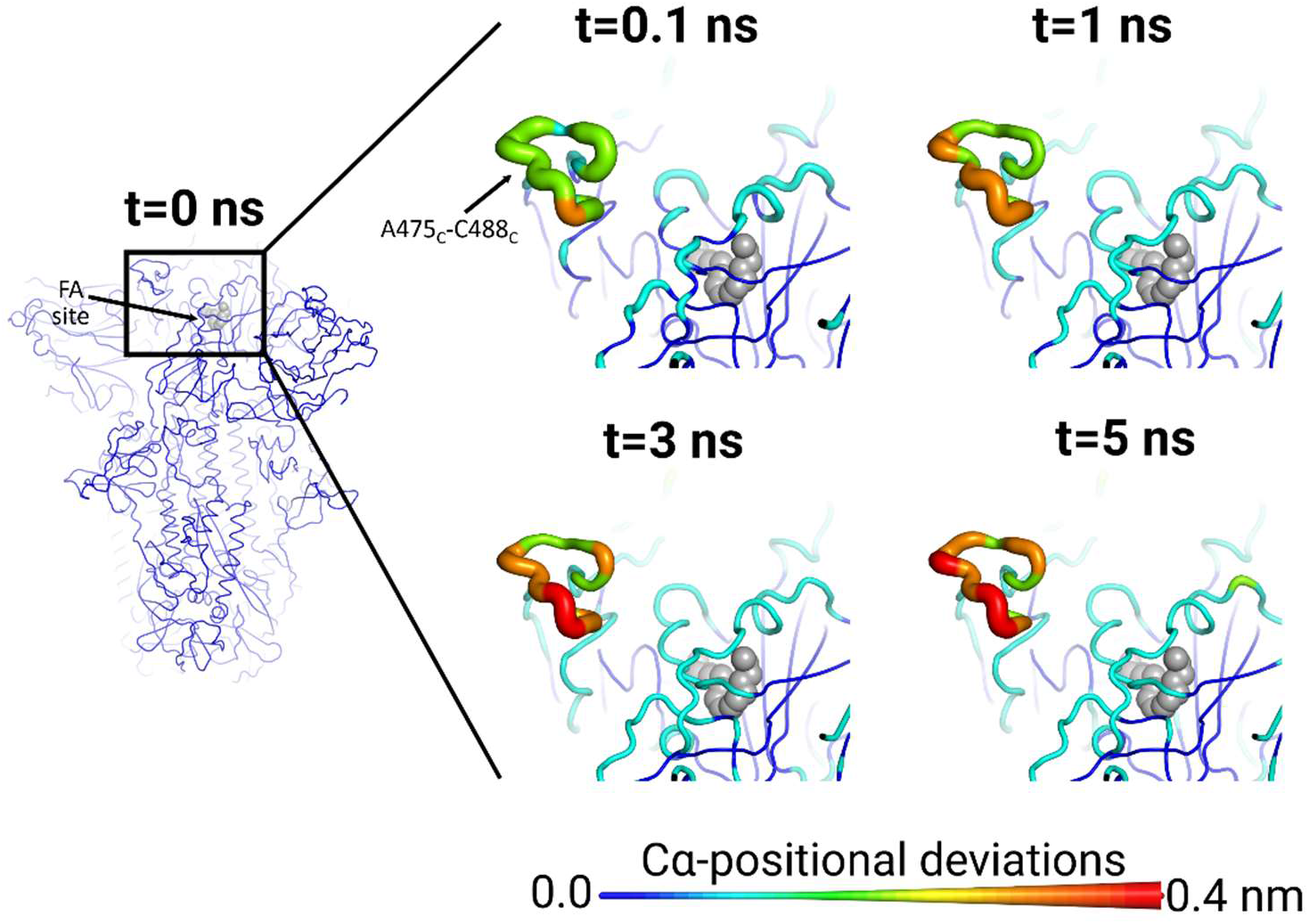
FA site allosterically affects the RBM in the SARS-CoV-2 spike. Average Cα-positional deviation around the first FA site at times 0, 0.1, 1, 3 and 5 ns following LA removal from the FA binding pockets. The Cα deviations between the simulations with and without LA were determined for each residue and averaged over the 90 pairs of simulations (Figures 2 and S4). The Cα average deviations are mapped onto the structure used as starting point for the LA-bound equilibrium simulations. Both the structure colours, and the cartoon thickness, relate to the average Cα-positional deviation values. The LA molecule is represented with grey spheres. The subscript letters in the labels correspond to the monomer ID. This figure shows the response around the first FA binding site formed by monomer A and C. Results for the other sites show similar connections between the FA sites and the RBMs (see Figures S9-S10 for the responses observed for the other two FA sites).

The NTDs also show a fast and significant response to LA removal, in particular H146-E156 and L249-G257 (Figures 4 and S11-S12). The NTD of the spike is a surface-exposed domain structurally linked to the RBD of a neighbouring monomer (*17, 38*). Although directly coupled to the RBD, the NTD does not bind to ACE2 (*5, 6*) and its function in SARS-CoV-2 infection remains unclear. The spike NTDs of other related coronaviruses have been suggested to play a role in infection (*44-46*) and are known epitopes for neutralising antibodies (*46, 47*). Human antibodies targeting the NTD of the SARS-CoV-2 spike have been isolated from convalescent COVID-19 patients (e.g. (*47-49*)) and this region was shown to be a super-antigenic site (*50*). A cryo-EM structure of the complex between the spike and the 4A8 monoclonal antibody shows that the NTD loops L141-E156 and R246-A260 (two of the regions that show the largest responses to LA removal in Figures 4 and S11-S12) directly mediate the interaction between the proteins (*48*). Both of these loops are candidates for vaccine and therapeutic developments (*48*). The conformational changes in the H146-E156 and L249-G257 segments are further transmitted, over the following 5 ns, to other parts of the NTD, namely S71-R78, N122-N125 and F175-F186. The N122-N125 segment is a conserved NxxN sequence motif present in the NTD of spikes from several coronaviruses, and its function remains unknown (*51*). The F175-F186 region is located immediately before a recently identified epitope for human antibodies (*49*). The S71-R78 segment is part of the GTNGTKR insertion shared by the SARS-CoV-2 and bat-CoV RaTG13 spikes but not the SARS-CoV spike (*51*). This motif, which is also found in structural proteins of several other viruses, and proteins from other organisms, has been suggested to allow the SARS-CoV-2 spike to bind to other receptors besides ACE2 (*51*). The coupling identified here between the FA site and specific regions of the NTD is remarkable and highlights the complex allosteric connections within the spike, with distant sites apparently able to modulate the response of the NTDs.

**Figure 4.**
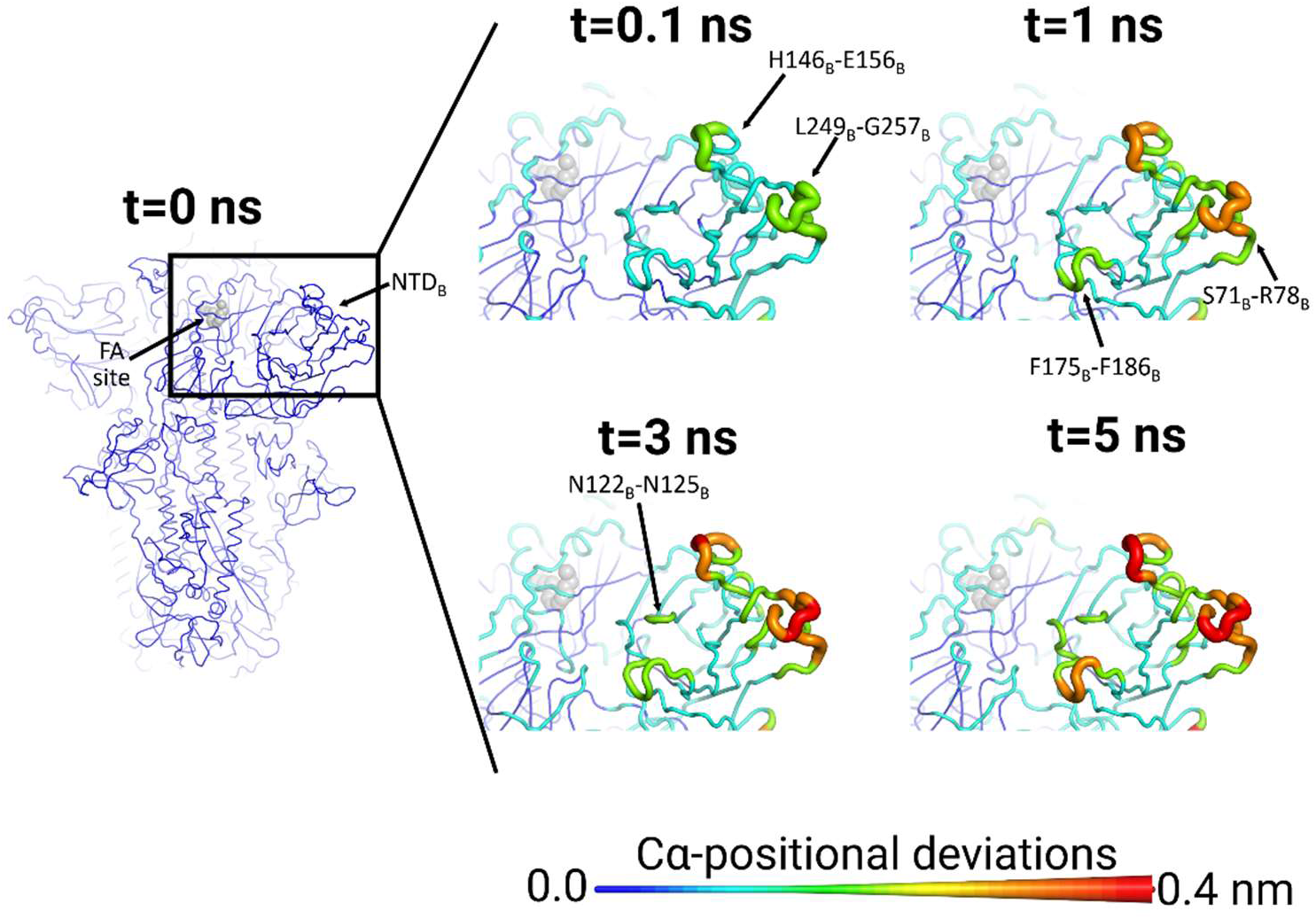
The FA site allosterically affects the NTD in the SARS-CoV-2 spike. Average Cα-positional deviation around the first FA site at times 0, 0.1, 1, 3 and 5 ns following LA removal from the FA binding pockets (see Figures S11-S12 for the responses observed for the other two FA sites). For more details, see the legend of Figure 3. This figure shows the response around the first FA binding site formed by monomer A and C. Similar results are observed for the other two FA sites and the NTD (Figures S11-S12).

Both the furin cleavage site and V622-L629 region, which are more than 40 Å away from the FA site, respond notably to the removal of LA (Figures 5 and S13-S14). Both regions respond rapidly, with a significant conformational response observed almost immediately after LA removal. The furin site is located at the boundary between the S1 and S2 subunits (*16, 17*) and furin cleavage is thought to be important for the activation of the spike (*19*). This site contains a polybasic PRRA insertion not found in other SARS-CoV-related coronaviruses (*52*). Cell-based assays show that deletion of the PRRA motif affects virus infectivity (*19, 52-56*). Note that in the simulations presented here, the furin site (located between R685 and S686) is not cleaved (see Supplementary Material).

**Figure 5.**
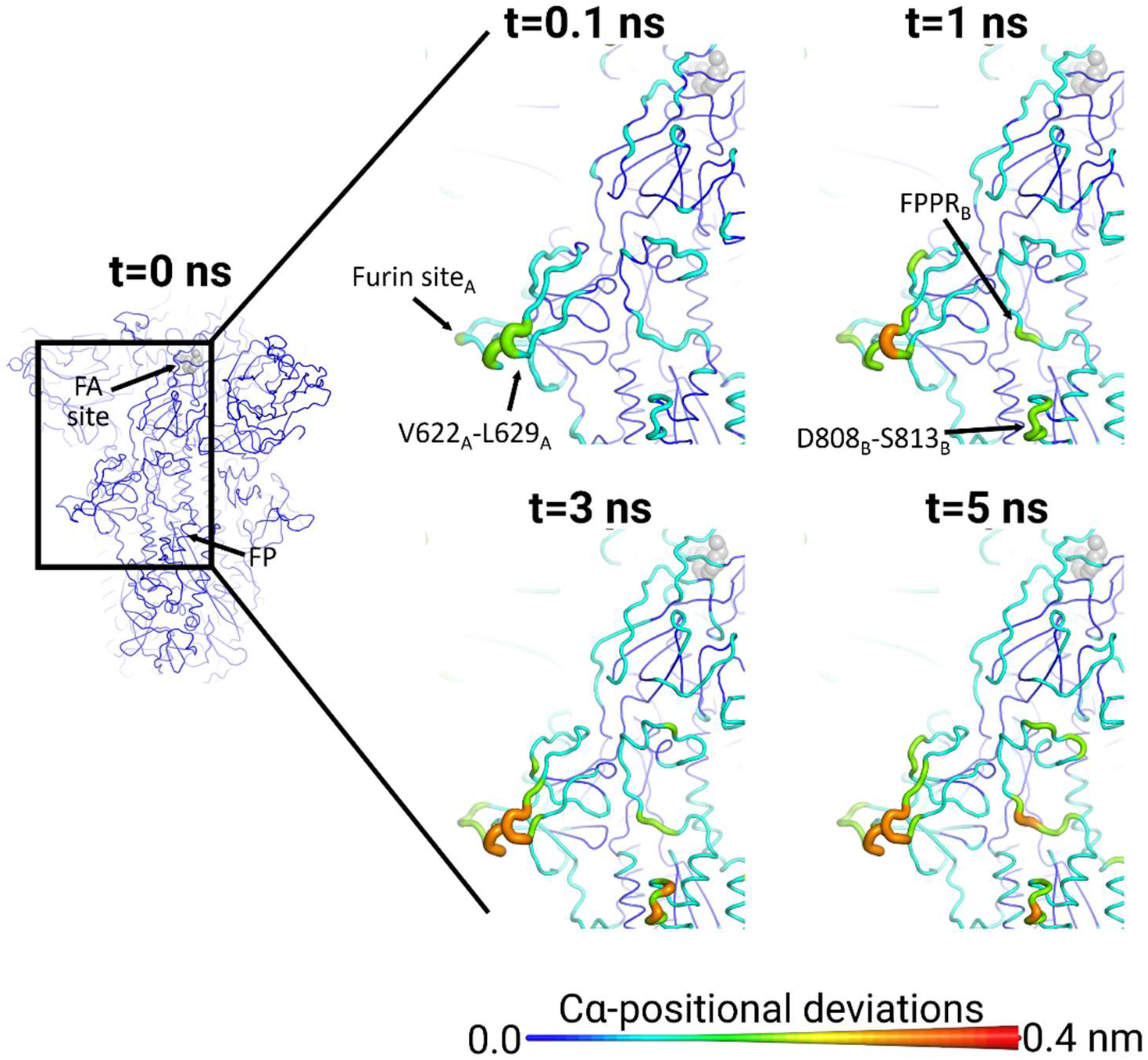
The FA site allosterically affects the furin site, FPPR, and the residues immediately preceding the S2’ cleavage site in the SARS-CoV-2 spike. Average Cα-positional deviations around the first FA site at times 0, 0.1, 1, 3 and 5 ns following LA removal from the FA binding pockets. This figure shows the response around the first FA binding site. A similar response is observed for all three monomers (see Figures S13-S14 for the responses observed for the other two FA sites). Analogous connections between the other two FA sites and the furin sites, FPPR and S2’ cleavage sites are also observed (Figures S13-S14). For more details, see the legend of Figure 3.

The furin site and V622-L629 regions are among the spike regions most affected by LA removal and show increasingly large deviations (larger than most other loop regions of the protein) over the simulations. The conformational changes in these regions propagate to segments immediately adjacent to the fusion peptide, namely the downstream FPPR and the upstream D808-S813. The FPPR is a ∼25-residue segment located in S2 immediately downstream of the fusion peptide which has been suggested to play an essential role in the structural transitions between pre- and post-fusion conformations of the spike (*18*). The D808-S813 region is located upstream of the fusion peptide, immediately preceding the S2’ protease recognition and cleavage site (R815) (*54*). Both proteolytic sites in the SARS-CoV-2 spike are known epitopes for neutralising antibodies (*57, 58*).

The close connection between the furin site, V622-L629, and the regions adjacent to the FP, identified here for the intact, wild-type spike is remarkable. Due to this crosstalk, mutations in or close to the furin site or V622-L629 are likely to affect signal transmission to the FP-surrounding regions, i.e. the FPPR and the S2’ cleavage site. This is worthy of experimental investigation.

### Dynamic response of the D614G spike

We also performed equilibrium and dynamical-nonequilibrium simulations of the D614G mutant spike. The D614G mutation is now dominant in SARS-CoV-2 lineages circulating worldwide (*59*) and confers increased transmissibility. Other variants are emerging as increasing numbers of infections provide further opportunities for mutations to arise. The B.1.1.7 variant, largely responsible for the surge in cases in the UK in the winter of 2020/21, has increased infectivity without the D614G mutation (*60*). However, three of the four variants involved in the April 2021 surge of cases in India do include D614G among the mutations (COVID-19 Genomics UK Consortium, https://www.cogconsortium.uk). That a single amino acid replacement of the aspartate residue in position 614 by a glycine can lead to more efficient viral transduction into host cells affording greater infectivity (*61-64*) is worthy of mechanistic exploration. Here, three equilibrium 200 ns MD simulations were performed for the locked form of the unglycosylated and uncleaved ectodomain region of the D614G spike modelled from a wild-type model (*17, 20*) based on the cryo-EM structure 6ZB5 (*17*) with and without LA bound (see Supplementary Material). In the original wild-type spike, D614 is located at the interface between monomers, with its sidechain directly interacting with residues across the subunit interface (*55*). RMSF profiles for the wild-type and D614G apo spikes are similar (Figure S15). However, unlike the wild-type with or without LA, one replicate of the D614G with LA bound exhibits enhanced fluctuation in the middle of the RBM corresponding to exposed loop residues Q474-N487 (Figure S15). Though the RBD in the closed conformation remains inaccessible for binding to ACE2, residues Q474-N487 of the RBM (shown in magenta in the insert in Figure S15F) may still provide a target for neutralizing antibodies in the closed conformation, depending on the degree of glycan shielding (*65*). Our equilibrium MD simulations of the unglycosylated, uncleaved wild-type and D614G LA-bound spikes suggest that the D614G may enhance RBM mobility. It is possible that an increase in flexibility in the RBM could influence the efficiency of glycosylation in this region (*66*), which could either increase or decrease the degree of glycan shielding, affecting epitope recognition.

The trans-interface interactions of the carboxylate of D614 in the wild-type systems involve four potential candidate residues, K854, K835, Q836 and T859. In our simulations, T859 came within 5 Å of D614 across an interface and made transient, weak interactions throughout the simulations of the apo and LA-bound spike systems (Figure S16). The carboxylate of D614 and NZ of K854 remains within salt-bridging distance throughout the wild-type simulations (Figure S17). The D614-K854 trans-subunit interface interaction dominates throughout the simulation time, regardless of the presence of bound LA (Figure S16). In our 200 ns open wild-type apo spike simulations, the trans-subunit interaction between D614 and K854 persisted for 99% of the frames across eight of the nine subunit interfaces. Nonetheless, this contact was lost at only one interface and that was between an open and closed chain in one of the repeats (Figure S16).

An analogous analysis was performed on the D614G mutant to establish whether K854 makes alternative hydrogen-bond or salt-bridge contacts across the 3 subunit interfaces (averaged over 3 × 200 ns replicates) in the absence of a partnering D614 carboxylate. In the D614G spike, K854 fails to find any alternative salt-bridge and only occasionally comes within hydrogen-bonding distance of residues Q613 and N317 (Figure S18). This supports the inference drawn from cryo-EM structures of the head region of the D614G spike that this mutation disrupts the inter-monomer salt-bridge and hydrogen bond networks in this region, which may cause reduced stability of the trimer. This corresponds to the observation that the D614G mutant was mostly in an open conformation on the EM grids and suggests that loss of the D614-K854 interaction somehow destabilises the closed conformation (e.g. (*67, 68*)).

Dynamical-nonequilibrium simulations for the D614G variant were also performed to test whether the D614G mutation affects the response of the spike to LA. These simulations used as a starting point the distribution of conformations taken from the equilibrium simulations of the locked form of the unglycosylated and uncleaved D614G spike with LA bound. The same perturbation as for the wild-type spike was applied to the system, namely LA removal. The Kubo-Onsager approach (*29-31*) was again used to extract the response of the system (Figure S19) and determine the statistical significance of the observed responses (Figure S20). In the D614G variant, there is notably less symmetry across the monomers in the response of the spike to LA removal, compared to the wild-type (Figures 6 and S21-S28). For instance, the amplitude of the structural response of the V266-L629 and furin site regions in monomer C (Figure S27) of the D614G is substantially smaller than in monomers B and A (Figures 6 and S28).

**Figure 6.**
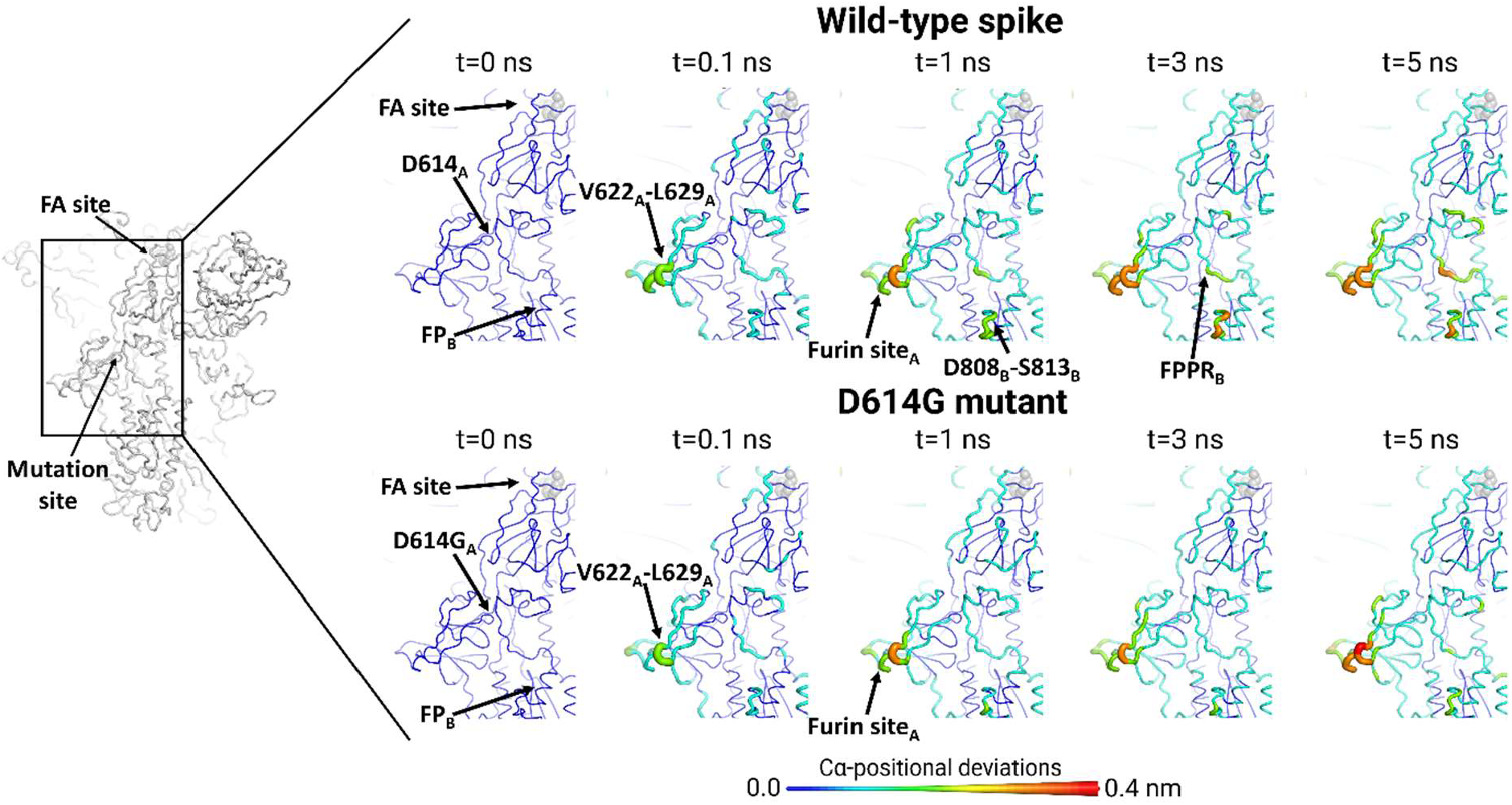
Response of the wild-type and D614G spike to LA removal. Average Cα-positional deviations around the first FA site at times 0, 0.1, 1, 3 and 5 ns following LA removal from the FA binding pockets (see Figures S27-S28 for the responses observed for the other two FA sites). The Cα deviations between the simulations with and without LA were determined for each residue, and the values averaged over the 90 pairs of simulations (Figures 2, S4 and S19). The structure colours, and cartoon thickness, relate to the average Cα-positional deviations. The LA molecule is represented with grey spheres. The subscript letters in the labels correspond to the monomer ID.

The conformational responses of the wild-type and D614G spikes can be directly compared because the same perturbation was used for both in the dynamical-nonequilibrium simulations. The conformational response of the NTDs and RBDs to LA removal is generally similar in the wild-type and D614G spike (Figures S21-S26) with small variations in the amplitude of the structural rearrangements of some functional motifs, e.g. RBMs. However, the D614G mutation significantly affects inter-monomer communication, with reduction of signal transmission from the furin site and V622-L629 of one monomer to the FPPR of another (Figures 6 and S27-S28) compared to the wild-type (Figures 5 and S13-S14). In the D614G spike, only minor deviations of the FPPR are observed. Furthermore, the region located upstream of the fusion peptide, namely D808-S813, also shows different rates of signal propagation between the wild-type and D614G proteins (Figures 6 and S27-S28). The differences identified here may relate to functionally important differences between the wild-type and D614G spikes. The results here show that the D614G mutation alters the allosteric networks connecting the FA site to the regions surrounding the FP, particularly the FPPR. There is reduced communication between the monomers in the D614G spike. As noted above, the response of the D614G spike to LA is also less symmetrical than the wild-type.

## Conclusions

Our findings show that changes in ligand occupancy at the FA site influence the dynamic behaviour of functionally important motifs distant from the FA site. The simulations identify a complex network of structural pathways connecting the FA sites to key structural motifs within the SARS-CoV-2 spike. These networks extend far beyond the regions surrounding the FA sites, with structural responses being observed in the RBM, NTD, furin site and FP-adjacent regions (Movie 1). The results also show strong crosstalk between the furin site, V622-L629 and the regions adjacent to the FP. Disrupting or altering these communication networks may be a novel strategy for drug development against COVID-19.

Simulations of the D614G spike show that this mutation affects communication between the FA site and the FPPR and the S2’ cleavage site. The D614G mutant shows reduced response of the FPPR and also a slower rate of signal propagation to the S2’ cleavage site when compared to the wild-type protein (Movie 2). These results indicate that the D614G mutation affects the allosteric behaviour and the response to linoleic acid of the spike, which may be related to the changes in viral fitness associated with this mutation (*69*).

The results here further highlight the potential of dynamical-nonequilibrium simulations for identifying pathways of allosteric communication (*32-34*) and suggest that this approach may be useful in analysing mutations and differences in functionally important dynamical behaviour, and possibly different effects of linoleic acid, between SARS-CoV-2 spike variants of clinical relevance.

## Acknowledgements

AJM and ASFO thank EPSRC (grant number EP/M022609/1) and BBSRC (grant number BB/R016445/1) for support. MD simulations were carried out using the computational facilities of the Advanced Computing Research Centre, University of Bristol (http://www.bris.ac.uk/acrc) under an award for COVID-19 research. We thank BrisSynBio, a BBSRC/EPSRC Synthetic Biology Research Centre (Grant Number:BB/L01386X/1) for funding DKS and ASFO and the BlueGem HPC system and EPSRC via HECBIOSIM (hecbiosim.ac.uk) for providing ARCHER/ARCHER2 time through a COVID-19 rapid response call. We also thank Oracle Research for Oracle Public Cloud Infrastructure (https://cloud.oracle.com/en_US/iaas) time under an award for COVID-19 research, the <u>Bristol UNCOVER Group</u> and the University of Bristol, for their support. I.B. acknowledges support from the EPSRC Future Vaccine Manufacturing and Research Hub (EP/R013764/1). C.S. and I.B. are Investigators of the Wellcome Trust (210701/Z/18/Z; 106115/Z/14/Z).

## Author contributions

Conceptualisation/design of the work: ASFO, DKS and AJM; Acquisition and analysis of the data: ASFO, DKS and AAI; Writing of the manuscript: ASFO, DKS and AJM; Review & Editing of the manuscript: ASFO, DKS, AAI, ADD, IB and CS; Funding Acquisition: AJM.

