## Supplementary Material for "The fatty acid site is coupled to functional motifs in the SARS-CoV-2 spike protein and modulates spike allosteric behaviour"

#### Materials and methods

##### Equilibrium simulations

The locked (three closed RBDs) and open (one raised RBD) wild-type spike proteins were modelled on 6ZB5 and the model from the map EMD-1146, respectively, as described in (*1, 2*). The wild-type sequence was checked by an EBI Blast (*3*) search, which revealed a 100% sequence identity to UNIPROT entry P0DTC2 for the wild-type SARS-CoV2 spike protein. The D614G mutant SARS2-CoV2 spike construct sequence was verified by alignment with Clustal Omega (*4*).

Loops were built using loop-builder in Chimera (UCSF) (*5*) and the integrity of the structures following loop builds was checked using PROCHECK (*6*) and iteratively improved by hand until the wild-type and D614G constructs had 99.5% and 99.6% residues in allowed regions of Ramachandran space respectively and main chain parameters were within (D614G exceeded) quality criteria for a structure resolved to 1.5 Angstroms (*6*). The locked constructs contained 42 disulphide bonds per trimer (Table S1). The chain with a raised RBD in the open constructs featured a N-terminal stretch that started at residue 15 and incorporated a disulphide between C15 and C136. Disulphide bonds play an important structural role in the spike, and so their inclusion is important.

**Table S1**. List of disulphide bonds present in each monomer in the open and locked wild-type and D614G mutant SARS2-CoV-2 spike models.

| |
|---|
| C15-C136 (raised RBD only) |
| C131-C166 |
| C291-C301 |
| C336-C361 |
| C379-C432 |
| C391-C525 |
| C480-C488 |
| C538-C590 |
| C617-C649 |

| |
|---|
| C662-C671 |
| C738-C760 |
| C743-C749 |
| C840-C851 |
| C1032-C1043 |
| C1082-C1126 |

The D614G structure was prepared with linoleate (LA) molecules bound in the same way as described for the wild-type system described previously in (*1, 2*). All wild-type and D614G structures contain no glycans and are uncleaved at the furin protease recognition site. Overall, six systems were prepared: the closed-apo wild-type and D614G spikes; the closed, LA-bound wild-type and D614G spikes (here referred to as locked-LA); the open apo (single RBD raised) wild-type and D614G spikes. For each system, 3 equilibrium MD simulations of 200 ns each were performed. The D614G and wild-type LA-bound and apo forms were prepared using GROMACS version 2019.2 (*7*). The D614G system was run with GROMACS version 2019.1 and the wild-type with version 2019.4 (*7*). The simulation conditions were the same as in our previous studies (*1, 2*). All the systems were considered sufficiently equilibrated after 50 ns (Figure S1).

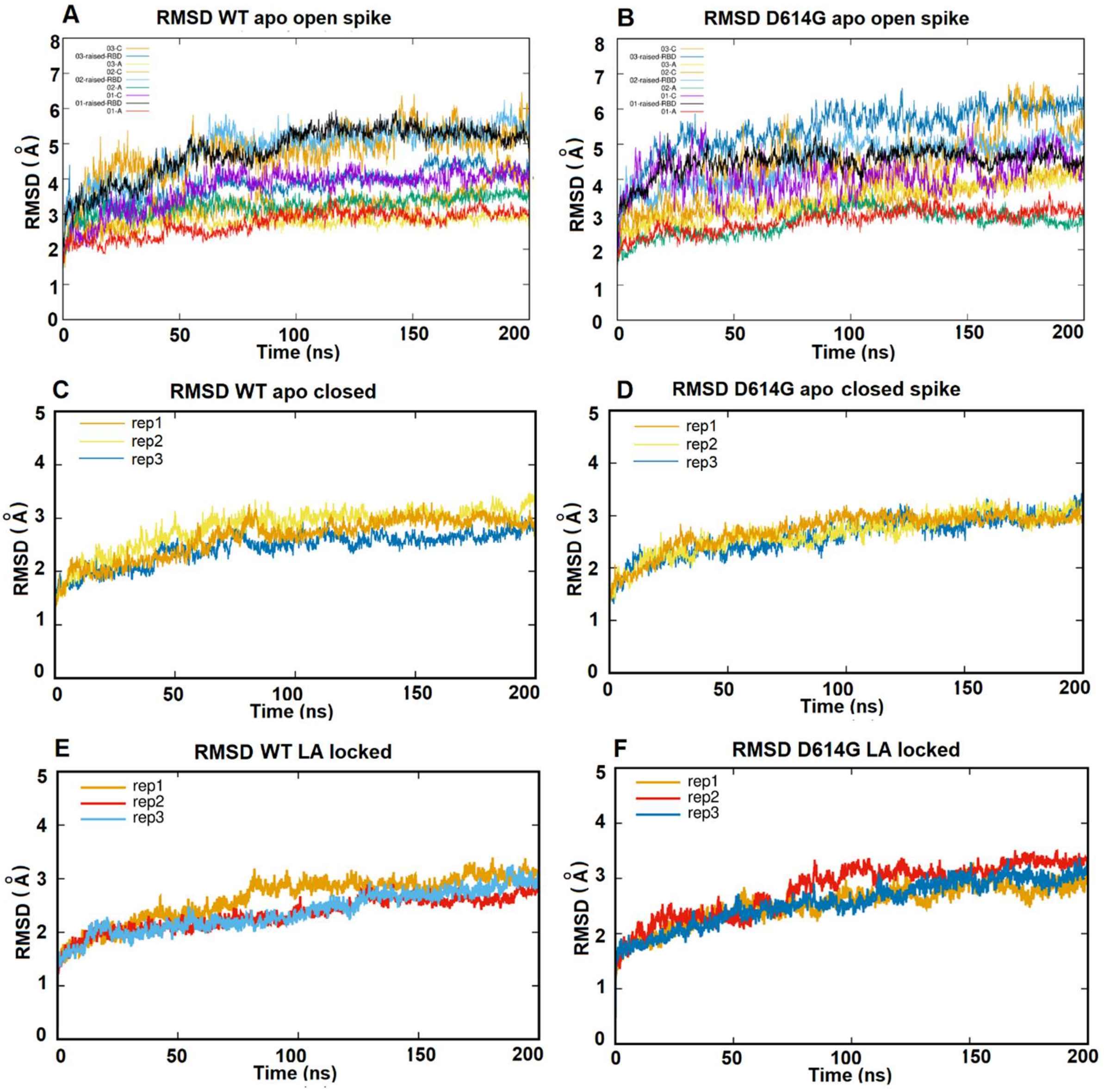


**Figure S1.** Root mean square deviation (RMSD) plots for the Cα positions from the replicate equilibrium MD simulations of the wild-type (WT) and D614G spikes. Panel (**A**) shows the RMSD for the open apo form (in which the RBD of chain B is raised) and (**B**) for the open apo form of the D614G spike. For the open spikes, RMSDs are shown for individual chains because they are in different conformations. RMSDs for the locked spike structures are averaged over the three chains of the trimer and are shown in (**C**) for the WT apo and (**D**) for the D614G mutant apo. Panel (**E**) shows RMSD from equilibrium MD simulations of the LA-bound locked WT and (**F**) for the D614G LA-bound locked spikes (each with 3 linoleates bound to each spike trimer).

#### Dynamical-nonequilibrium simulations

To study the structural changes associated with signal propagation within the unglycosylated, uncleaved wild-type and D614G mutant spike proteins, two sets of 90 short dynamical-nonequilibrium simulations each were carried out. Dynamical-nonequilibrium simulations have been applied successfully to various proteins, e.g. ABC transporters (*8, 9*), nicotinic acetylcholine receptors (*10, 11*), and class A β-lactamases (*12*). The Kubo-Onsager approach (*13-15*) is used to extract the conformational response of the proteins to ligand removal. In this approach (*13-15*), the response of a system to a perturbation is directly computed by calculating the difference of a given property between the simulations with and without the perturbation. Subtracting the perturbed and unperturbed pairs of simulations at a given time and averaging the results over many replicates allows not only the identification of the events associated with signal propagation, but also the determination of the statistical significance of the observations (*13-15*).

Here, the perturbation was generated by the (instantaneous) removal of the LA molecules from the fatty acid binding sites in the both the wild-type and D614G spike proteins from SARS-CoV-2. Note that the perturbation used here is not intended to represent the physical process of unbinding, but rather it was designed to drive a rapid response and force signal propagation within the protein, as its conformation adapts to LA removal. Furthermore, it should also be noted that these nonequilibrium simulations (due to their short timescales) are not attempting to sample state transitions (e.g. opening or closing of the RBDs). Instead, they allow the identification of the initial stages conformational changes associated with signal propagation within the protein. Nonetheless, the structural elements of the communication pathways identified here are likely to be involved in the response to LA binding/unbinding and the conformational rearrangements required for the transitions between states.

A graphical representation of the procedure used to set up the dynamical-nonequilibrium simulations is shown in the top panel of Figure S2. The starting conformations for the dynamical-nonequilibrium apo simulations were obtained from the equilibrated part of the equilibrium locked LA-bound simulations (see “equilibrium simulations” section above). Conformations were taken every 5 ns, and in each frame, the LA molecules were (instantaneously) removed from the FA pockets. The resulting apo system was then simulated for 5 ns (Figure S2). The simulation conditions for the nonequilibrium simulations were identical to the equilibrium ones described above. A total of 90 short apo dynamical-nonequilibrium simulations were performed (30 simulations per replicate) for each system. Note that to maintain the electroneutrality of the system (required for the PME (*16*) method), three positive ions were also removed from the solvent.

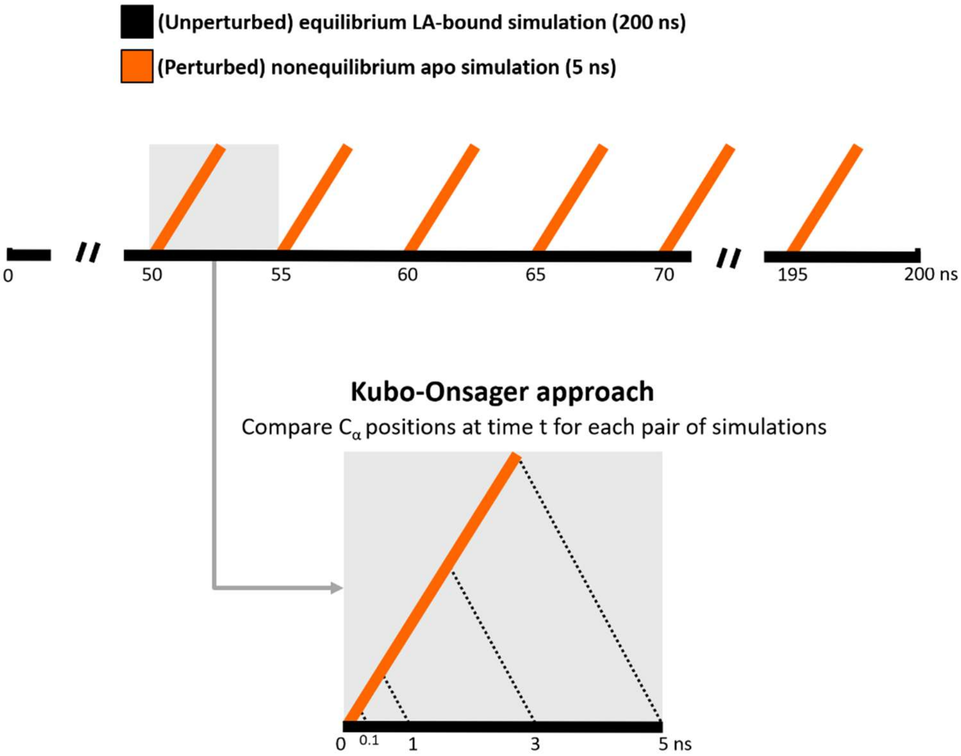


**Figure S2.** Schematic description of the procedure used to set up and analyse the dynamical-nonequilibrium simulations. For both the wild-type and D614G mutant, 3 equilibrium MD simulations, 200 ns each, were performed for the unglycosylated and uncleaved (no cleavage at the S1/S2 interface) locked spike. These equilibrium simulations (black lines) were, then, used to generate starting structures for the short apo dynamical-nonequilibrium simulations (orange lines). From the equilibrated part of each LA-bound simulation (from 50-200 ns), conformations were extracted every five nanoseconds, and the perturbation was introduced. Each short nonequilibrium apo simulation was simulated for 5 ns. The Kubo-Onsager (*13-15*) approach was used to extract the response of the system to LA annihilation from the FA pockets (bottom panel). For that, for each pair of unperturbed LA-bound and perturbed apo simulations, the positional deviations of each Cα at equivalent times (namely 0, 0.1, 1, 3 and 5 ns) were determined and averaged over all 90 simulations.

The Kubo-Onsager (*13-15*) approach was used to extract the response of the protein to LA removal (see the bottom panel in Figure S2). For each pair of unperturbed LA-bound equilibrium and perturbed apo nonequilibrium simulations, the difference in positions for each Cα was determined at equivalent points in time, namely after 0, 0.1, 1, 3 and 5 ns of simulation. The pairwise comparison between the positions of Cα atoms allows for the direct identification of the most important conformational rearrangements. We do not examine here the side chain responses. These would also be interesting, although will be more noisy. The Cα-positional deviations between the states at each point in time were averaged over all simulations. The statistical significance of the structural changes identified here is demonstrated by the low standard error of the averages (Figures S5 and S20).

### Supporting figures

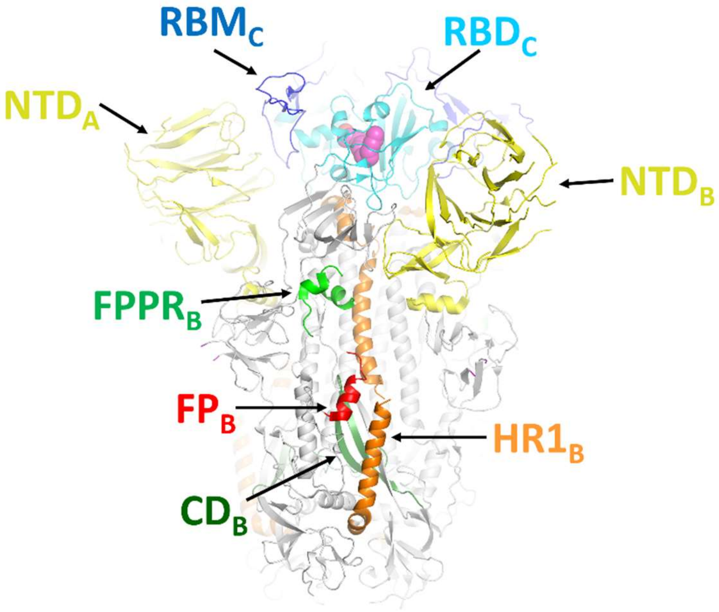


**Figure S3.** Cryo-EM structure of the ectodomain of the SARS-CoV-2 spike trimer (*2*) with some relevant structural motifs highlighted: N-terminal domain, NTD (residues 14-306)-yellow, receptor-binding domain, RBD (residues 331-528)- cyan; receptor binding motif, RBM (residues 438-506)- blue ; fusion peptide, FP (residue 816-827)- red; fusion-peptide proximal region, FPPR (residues 828-853)- green; heptad repeat 1, HR1 (residues 910-985)- orange; central helix, CH (residues 985-1035)- orange; connector domain, CD (residues 1035-1068)- dark green. Linoleic acid (LA) is shown with magenta spheres. Note that this structure represents the ectodomain of the spike trimer in the locked conformation (three closed RBDs).

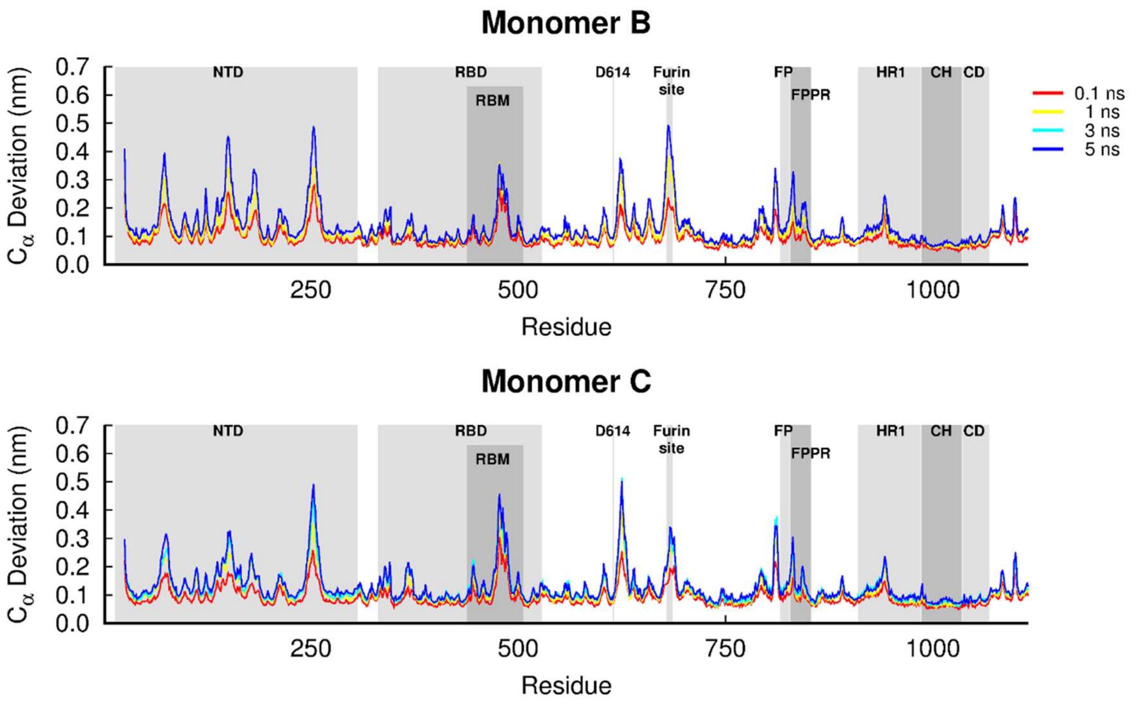


**Figure S4.** Average Cα-positional deviations for the second and third monomers in the five nanoseconds after LA removal from the FA sites in the wild-type spike protein from SARS-CoV-2. The average deviations were calculated using the Kubo-Onsager approach (*13-15*) for the pairwise comparison between the nonequilibrium apo and equilibrium LA-bound simulations. The positions of some important structural motifs are highlighted in grey, namely the N-terminal domain (NTD), receptor-binding domain (RBD), receptor-binding motif (RBM), fusion peptide (FP), fusion-peptide proximal region (FPPR), heptad repeat 1 (HR1), central helix (CH) and connector domain (CD).

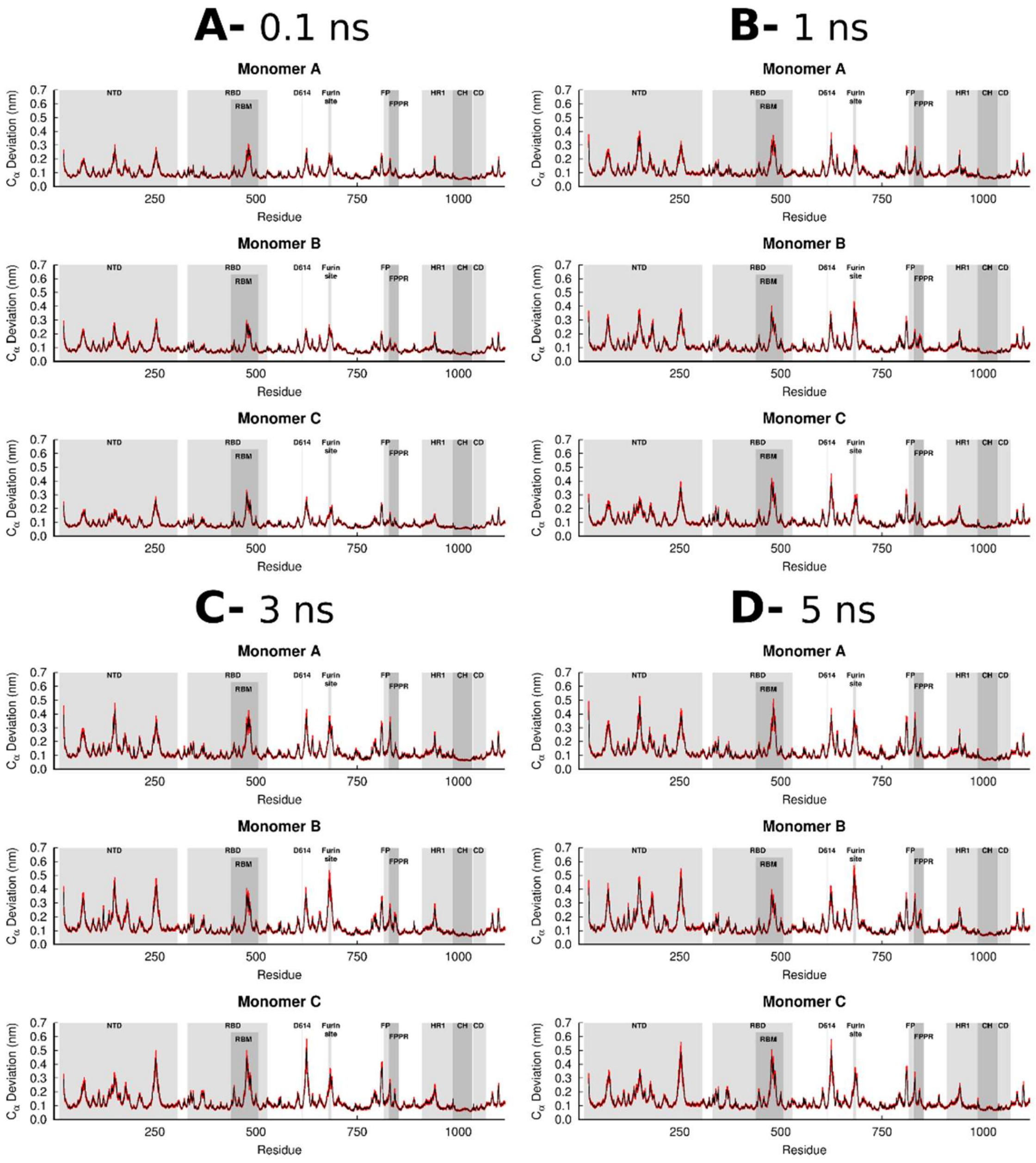


**Figure S5.** Average Cα-positional deviations (and corresponding standard errors) in the 0.1 (**A**), 1 (**B**), 3 (**C**) and 5 (**D**) ns after LA removal from the FA site in the wild-type spike protein. The average deviations were calculated using the Kubo-Onsager approach (*13-15*) for the pairwise comparison between the nonequilibrium apo and equilibrium LA-bound simulations. The vertical red lines represent the standard error of the mean. The positions of some important structural motifs are highlighted in grey, namely the N-terminal domain (NTD), receptor-binding domain (RBD), receptor-binding motif (RBM), fusion peptide (FP), fusion-peptide proximal region (FPPR), heptad repeat 1 (HR1), central helix (CH), connector domain (CD). Please zoom in to the image for detailed visualisation.

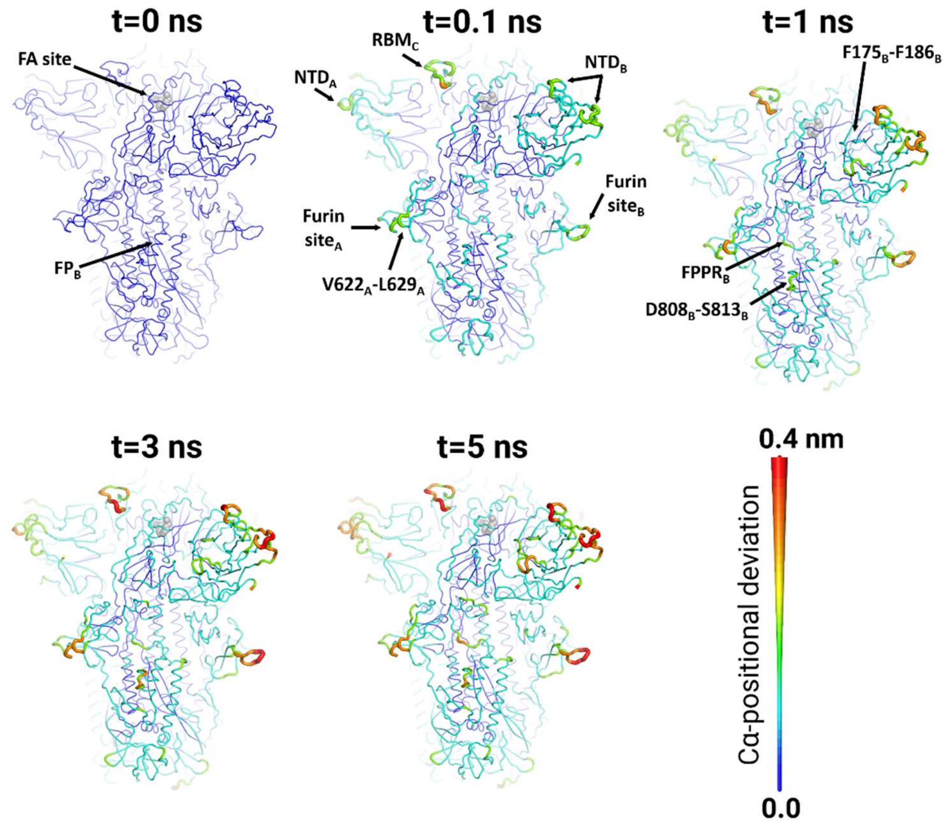


**Figure S6.** Mapping of the average Cα-positional deviations around the first FA site in the five nanoseconds following LA removal in the wild-type SARS-CoV-2 spike. The $C_\alpha$ deviations between the nonequilibrium apo and equilibrium LA-bound simulations at specific times (0, 0.1, 1, 3 and 5 ns) after LA removal were calculated as a function of the residue number. The final deviation values correspond to the average obtained over all 90 pairs of simulations (Figures 2 and S4). The Cα average deviations are mapped onto the structure used as the starting point for the LA-bound equilibrium simulations. Note that in this image, both structure colours and cartoon thickness relates to the average Cα-positional deviation values. A LA molecule is represented with grey spheres. The subscript letters in the labels correspond to the monomer ID. Please zoom in to the picture for detailed visualisation.

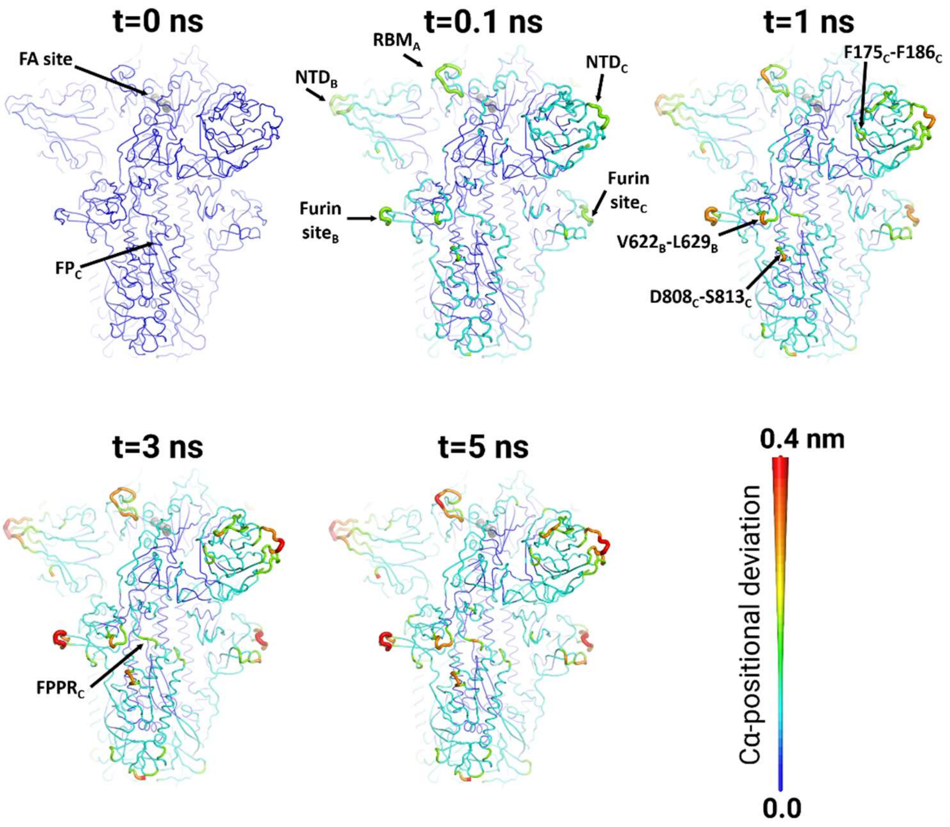


**Figure S7.** Mapping of the average Cα-positional deviations around the second FA site in the five nanoseconds following LA removal in the wild-type SARS-CoV-2 spike. The $C_\alpha$ deviation between the nonequilibrium apo and equilibrium LA-bound simulations at specific times (0, 0.1, 1, 3 and 5 ns) after LA removal were calculated as a function of the residue number. Please zoom in to the picture for detailed visualisation. For more details, see the legend of Figure S6.

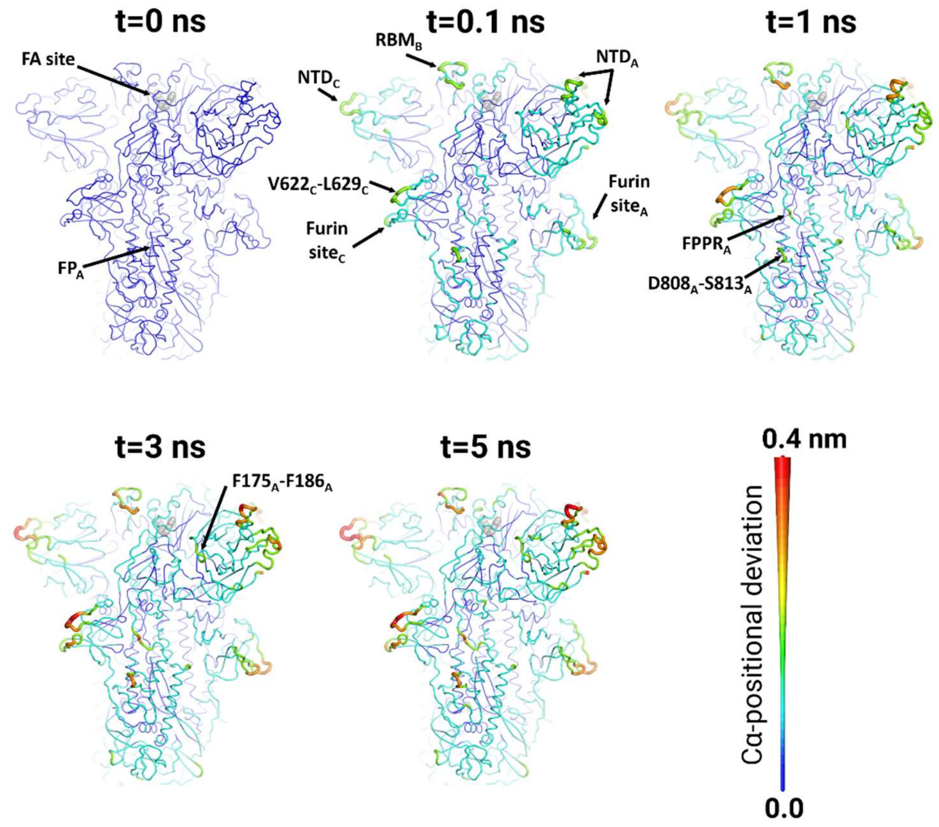


**Figure S8.** Mapping of the average Cα-positional deviations around the third FA site in the five nanoseconds following LA removal in the wild-type SARS-CoV-2 spike. The $C_\alpha$ deviation between the nonequilibrium apo and equilibrium LA-bound simulations at specific times (0, 0.1, 1, 3 and 5 ns) after LA removal were calculated as a function of the residue number. Please zoom in to the picture for detailed visualisation. For more details, see the legend of Figure S6.

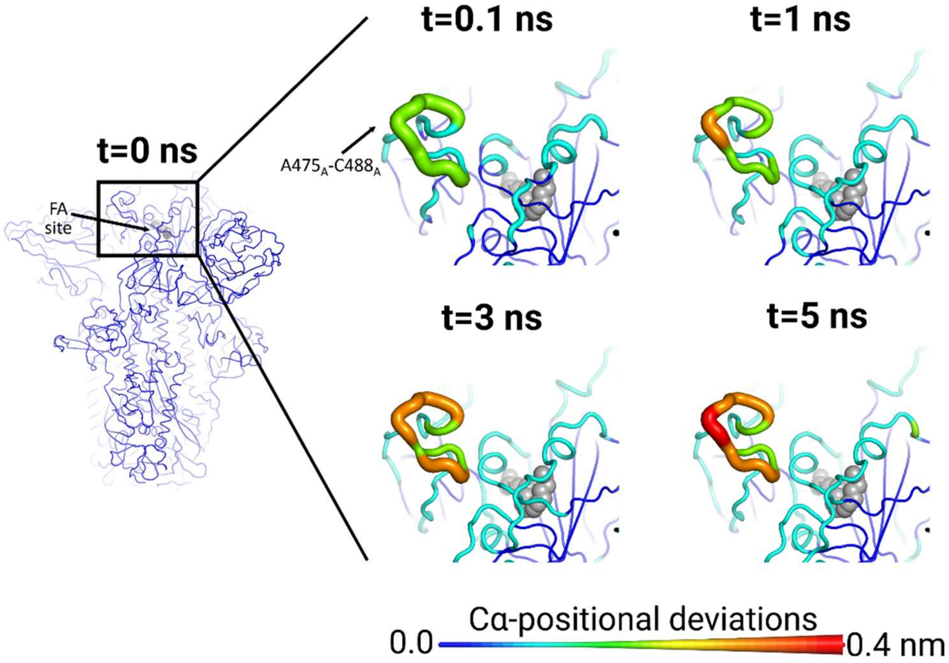


**Figure S9.** The second FA site allosterically affects the $RBM_A$ (residues 475-488) in the wild-type SARS-CoV-2 spike. Average Cα-positional deviation at times 0, 0.1, 1, 3 and 5 ns following LA removal from the FA binding pockets. The Cα deviations between the simulations with and without LA were determined for each residue, and the final values were averaged over the 90 pairs of simulations (Figures 2 and S4). The Cα average deviations are mapped onto the structure used as the starting point for the LA-bound simulations. Note that in this image, both structure colours and cartoon thickness relates to the average Cα-positional deviation values. LA molecule is represented with grey spheres. The subscript letters in the labels correspond to the monomer ID.

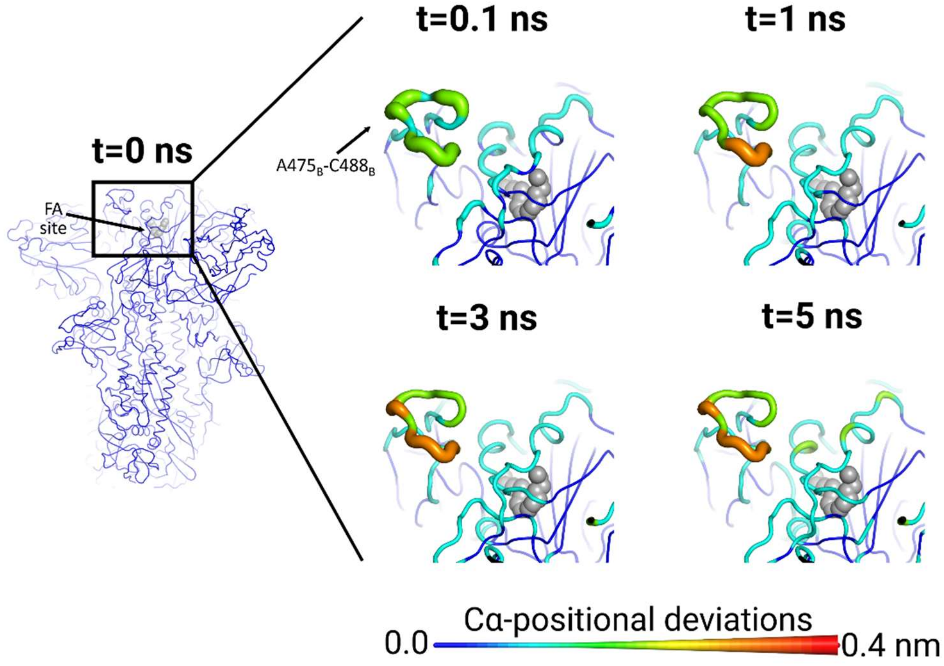


**Figure S10.** The third FA site allosterically affects the $RBM_B$ (residues 475-488) in the wild-type SARS-CoV-2 spike. Average Cα-positional deviation at times 0, 0.1, 1, 3 and 5 ns following LA removal from the FA binding pockets. For more details, see the legend of Figure S9.

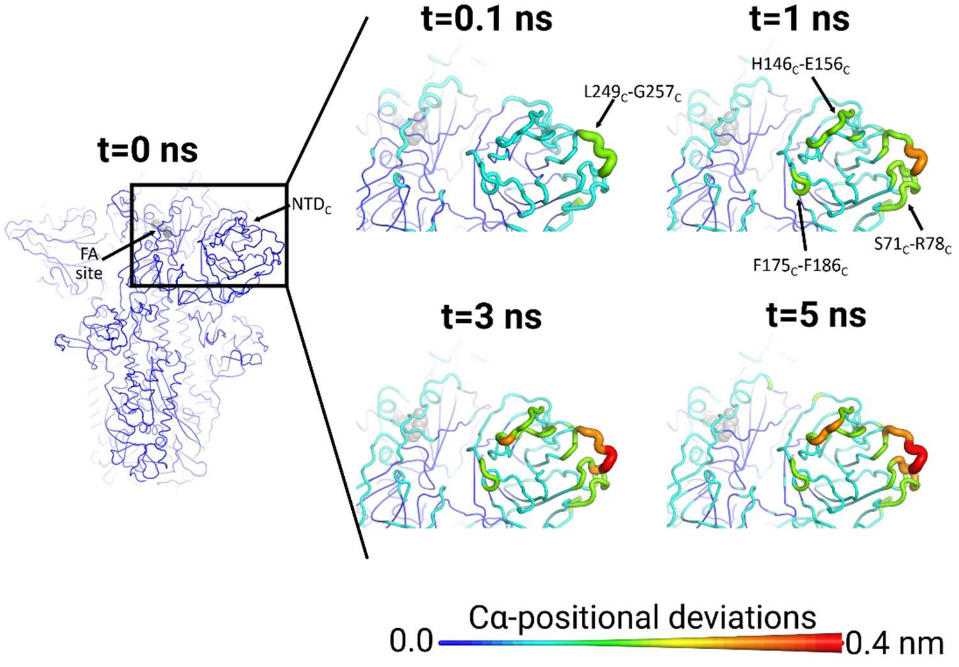


**Figure S11.** The second FA site allosterically affects the $NTD_C$ in the wild-type SARS-CoV-2 spike. Average Cα-positional deviation at times 0, 0.1, 1, 3 and 5 ns following LA removal from the FA binding pockets. For more details, see the legend of Figure S9.

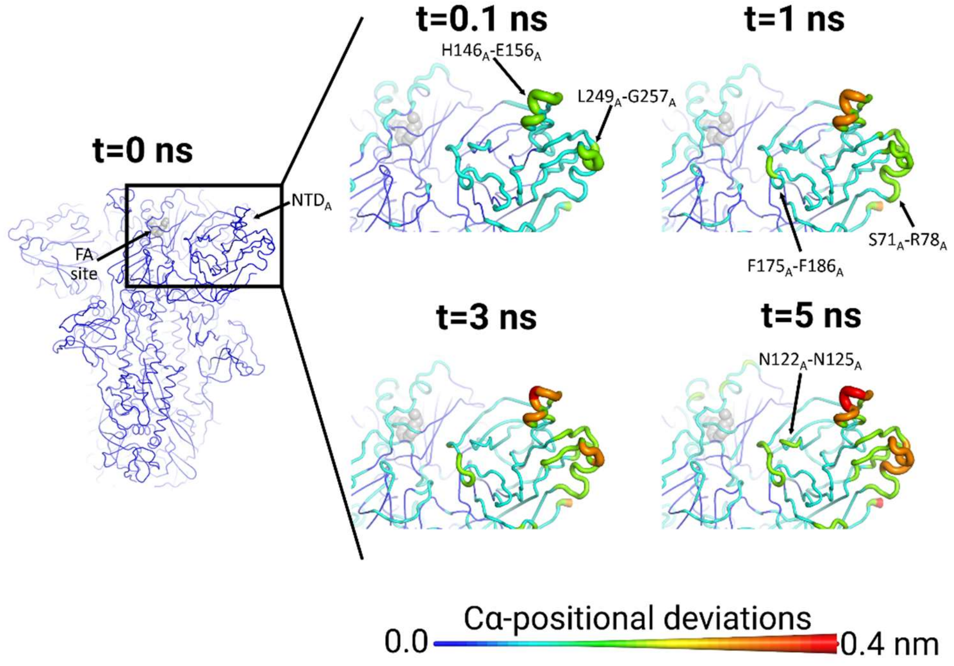


**Figure S12.** The third FA site allosterically affects the $NTD_A$ in the wild-type SARS-CoV-2 spike. Average Cα-positional deviation at times 0, 0.1, 1, 3 and 5 ns following LA removal from the FA binding pockets. For more details, see the legend of Figure S9.

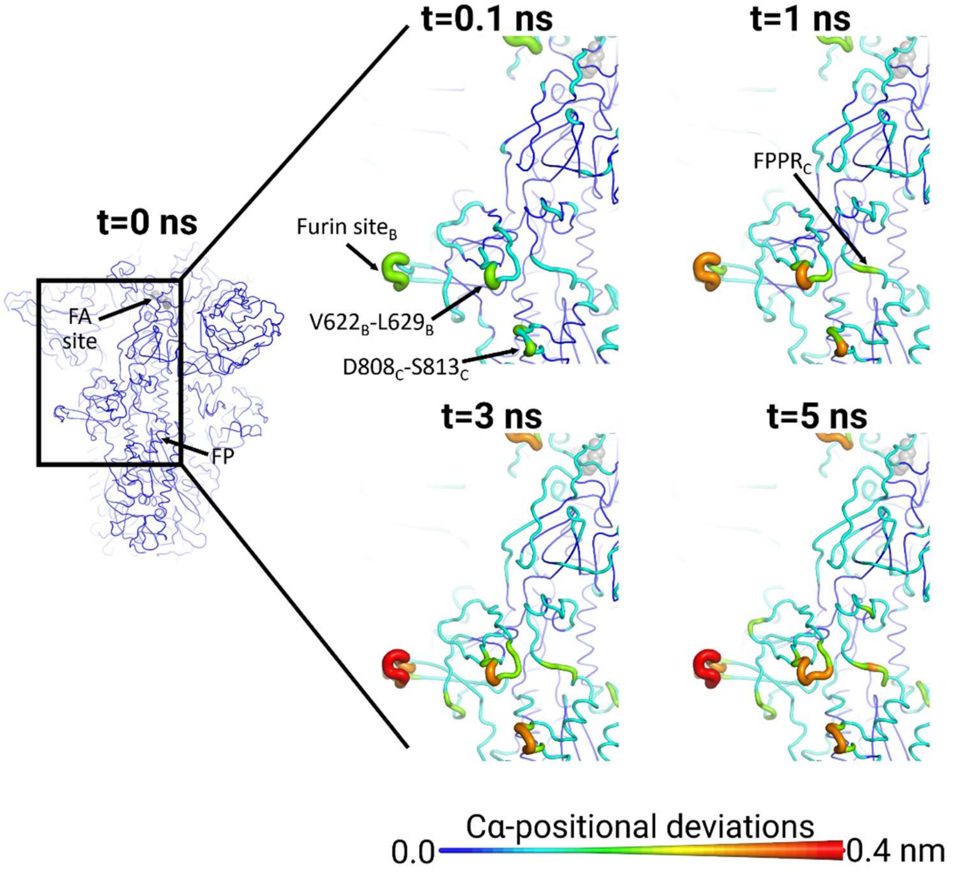

**Figure S13.** The second FA site allosterically affects the furin site$_B$, FPPR$_C$ and the residues immediately preceding the S2' cleavage site$_C$ in the wild-type SARS-CoV-2 spike. Average Cα-positional deviation at times 0, 0.1, 1, 3 and 5 ns following LA removal from the FA binding pockets. Note that the S2' protease recognition and cleavage site is located at position 815 (*17*). For more details, see the legend of Figure S9.

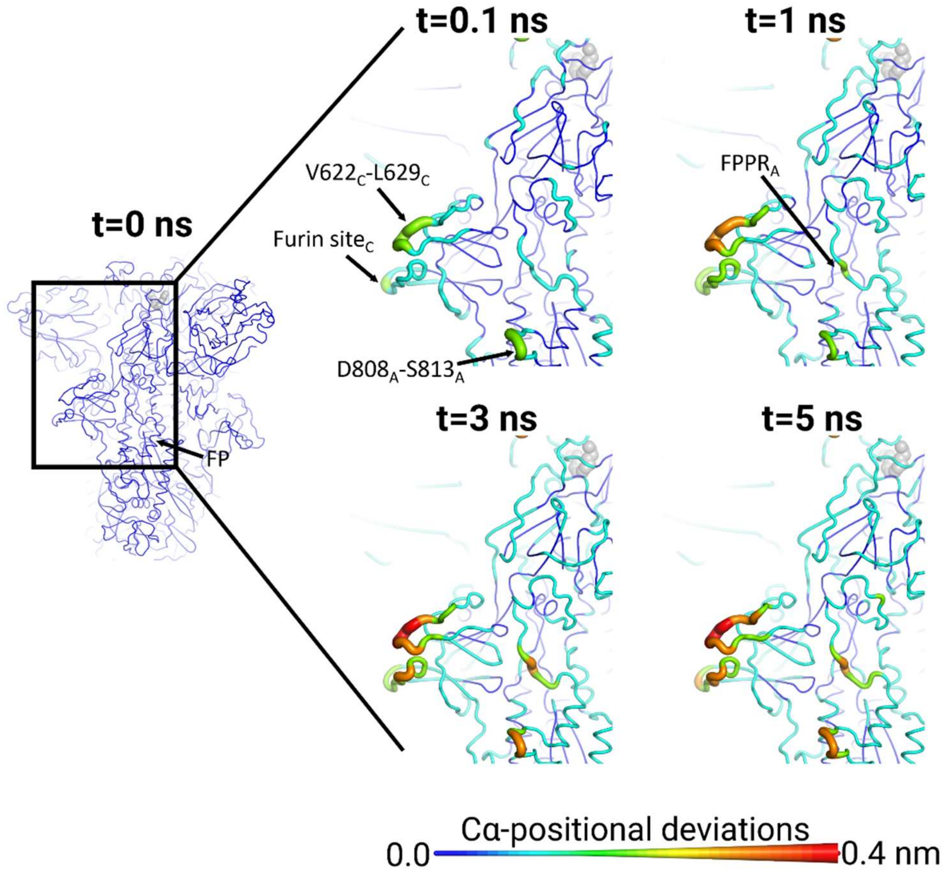


**Figure S14.** The third site allosterically affects the furin site$_C$, FPPR$_A$ and the residues preceding the S2' cleavage site$_A$ (R815) in the wild-type SARS-CoV-2 spike. Average Cα-positional deviation at times 0, 0.1, 1, 3 and 5 ns following LA removal from the FA binding pockets. Note that the S2' protease recognition and cleavage site is located at position 815 (*17*). For more details, see the legend of Figure S9.

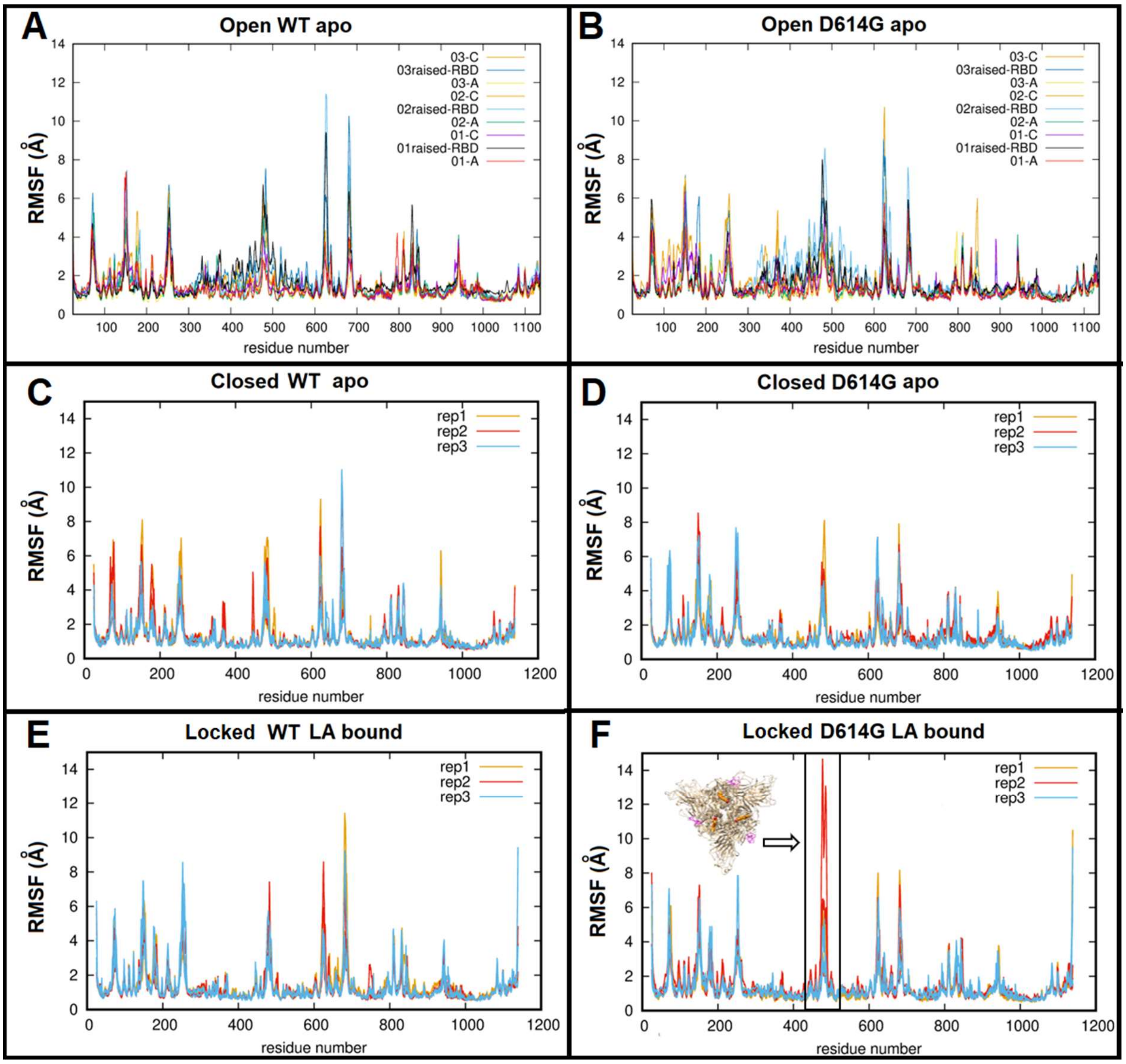


**Figure S15**. Root mean square fluctuations (RMSF) plots for the open wild-type apo (**A**), open D614G apo (**B**), closed wild-type apo (**C**), closed D614G apo (**D**), wild-type LA-locked (**E**) and for D614G LA-locked (**F**). The RMSF profiles were determined over the 200 ns of simulation shown as separate plots for the open structures as the chains were in non-identical conformations. The RMSF profiles for the locked structures were averaged over the three chains per trimer. Panel F shows that one of the D614G replicates with LA bound experienced enhanced fluctuations spanning in the exposed RBM. The insert shows the RBM residues in magenta that become more mobile in this replicate. Please zoom in to the picture for detailed visualisation.

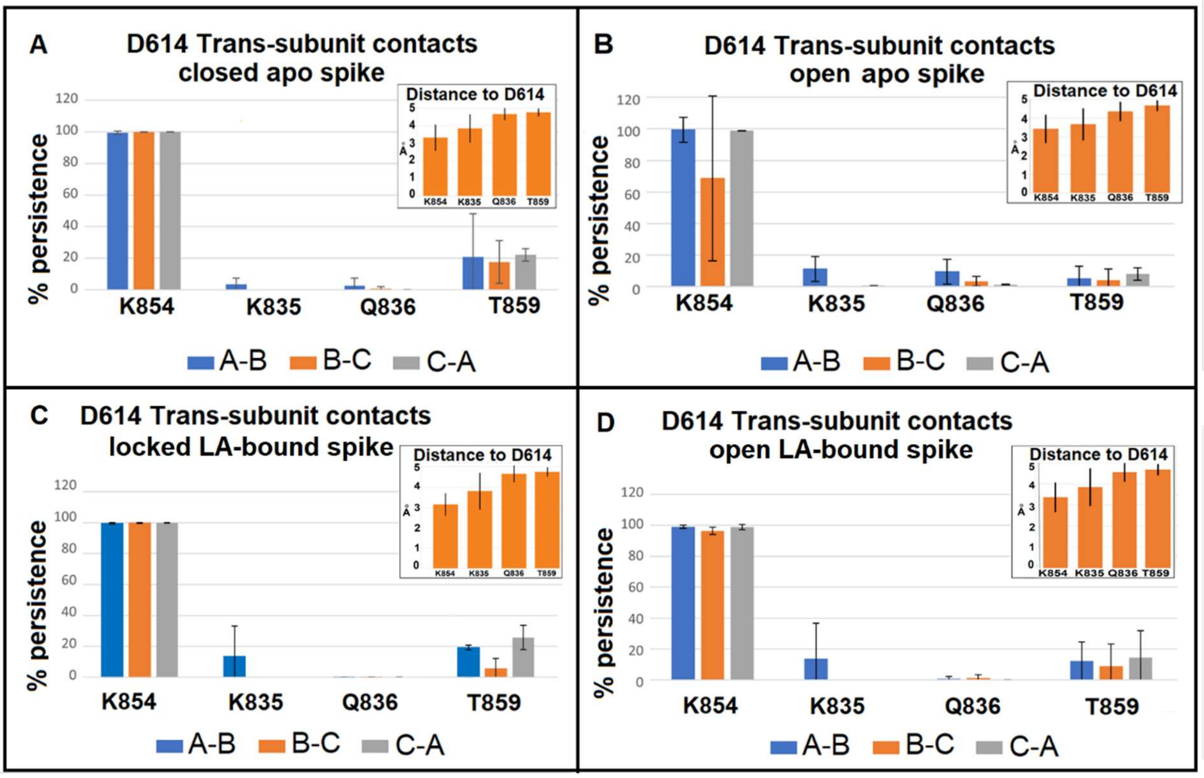


**Figure S16.** D614 interactions in the wild-type spike. Proportion of simulation time for which a 5 Å proximity cut-off was satisfied for the side chain carboxylate atoms of D614 to interact with polar atoms making inter-subunit interactions in the wild-type apo closed (**A**) and open (**B**) (in which the RBD of the B chain is raised) and the corresponding linoleate bound locked conformation (**C**) and open (**D**) simulations. Each system underwent three replicates, each of 200 ns. The inset histograms show the distance between the carboxylate atoms of D614 with the corresponding contacting residue atoms, averaged over time for all interfaces and all repeat simulations, with standard error bars. The generous 5 Å cut-off was chosen to extract contacts between D614 in the wild-type and any contacting inter-subunit residues. This distance was deemed appropriate to compensate for the resolution of the data (1000 frames were examined per 200 ns trajectory) in order to catch contacts that may have moved closer but may otherwise have been missed. Panels A, B, C and D show that in the wild-type, the D614 of the SARS-CoV-2 spike protein makes persistent (almost 100%) and short-range (average distance ~3.2 Å) interactions predominantly with K854 across each subunit interface, contributing to trimer cohesion. In summary, D614 interacts with K854 persistently across the trimer interface with only one exception when the contact between D614 and K854 was lost at a single subunit interface (this happened between the chain with a raised RBD and its neighbour at a B-C interface shown in panel B).

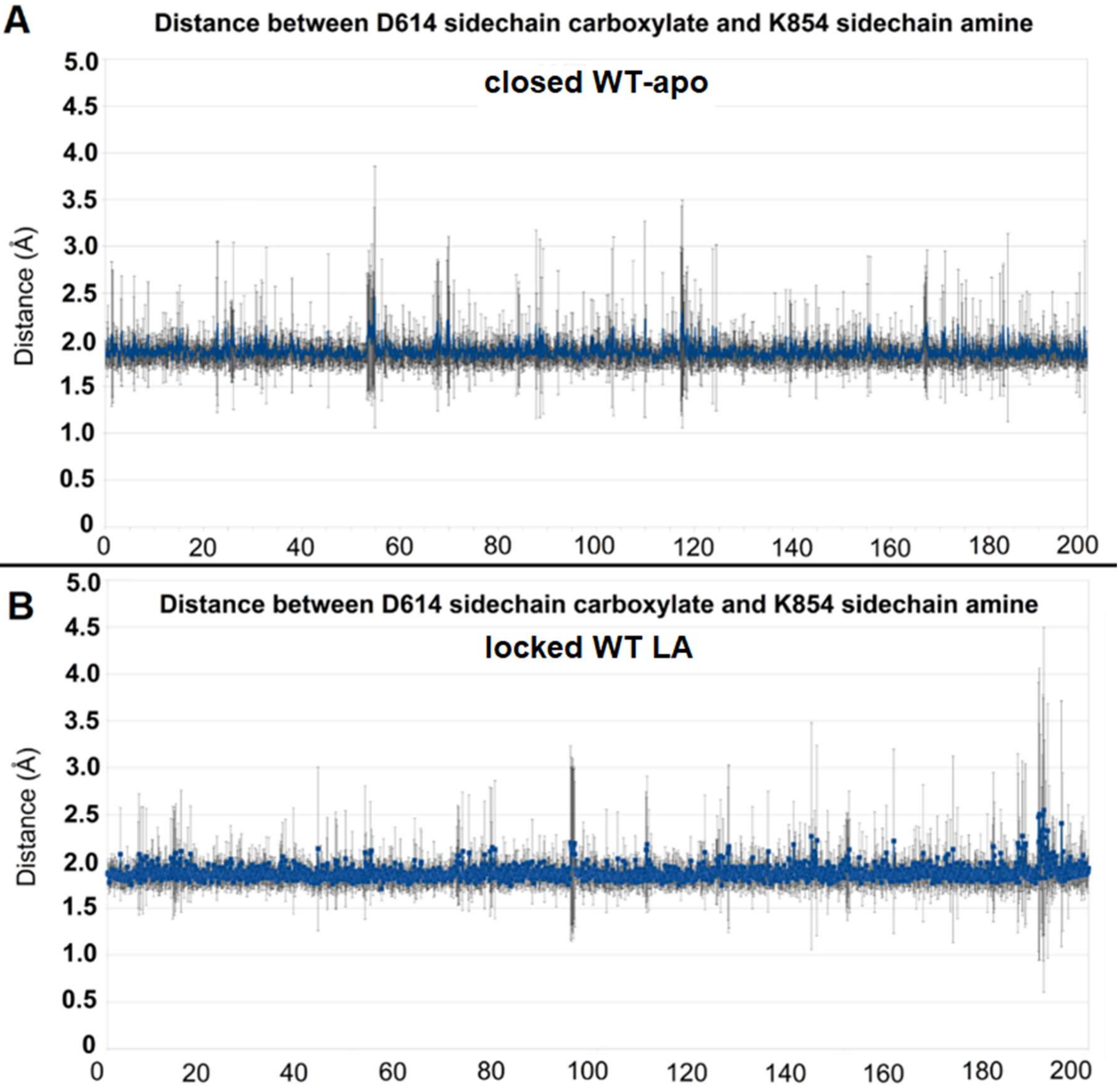


**Figure S17**. Distance between the carboxylate atoms of D614 and the sidechain amine nitrogen of the K854 on the neighbouring subunit in the closed wild-type apo (**A**) and the locked wild-type LA-bound (**B**) SARS-CoV-2 spike protein. The distances were averaged over the 3 subunit interfaces per trimer and over replicates. The vertical lines represent the standard deviation of the mean. In summary, the distance between D614 and K854 is unaffected by the presence of LA in the FA site under these simulation conditions.

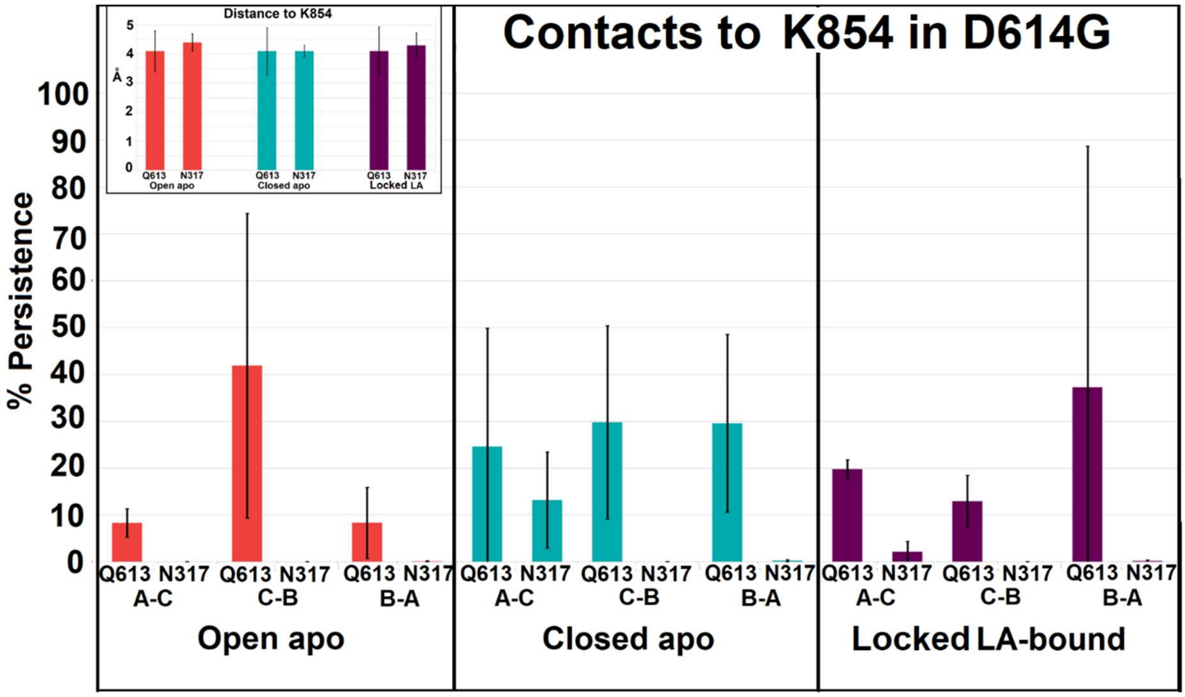


**Figure S18.** K854 contacts in the D614G mutant. In order to explore whether K854 was able to compensate for the loss of the salt bridge in the D614G mutant, contacts made by the side chain of K854 were examined over the course of 3 replicates of 200 ns each for the D614G mutant in the open apo and closed states with and without LA bound. This plot shows the proportion of simulation time for which a 5 Å proximity cut-off was satisfied for inter-subunit interactions of the terminal side chain amine nitrogen (NZ) atom of K854 with the polar side-chain atoms of inter-subunit (A-C, C-B, B-A) interacting residues. Inset histograms show the averaged distances between. K854 finds no compensating salt-bridges across the subunit interface and the two residues, Q613 and N317, capable of providing inter-subunit H-bonding did so transiently and with interactions averaging over 4 Å (see inset).

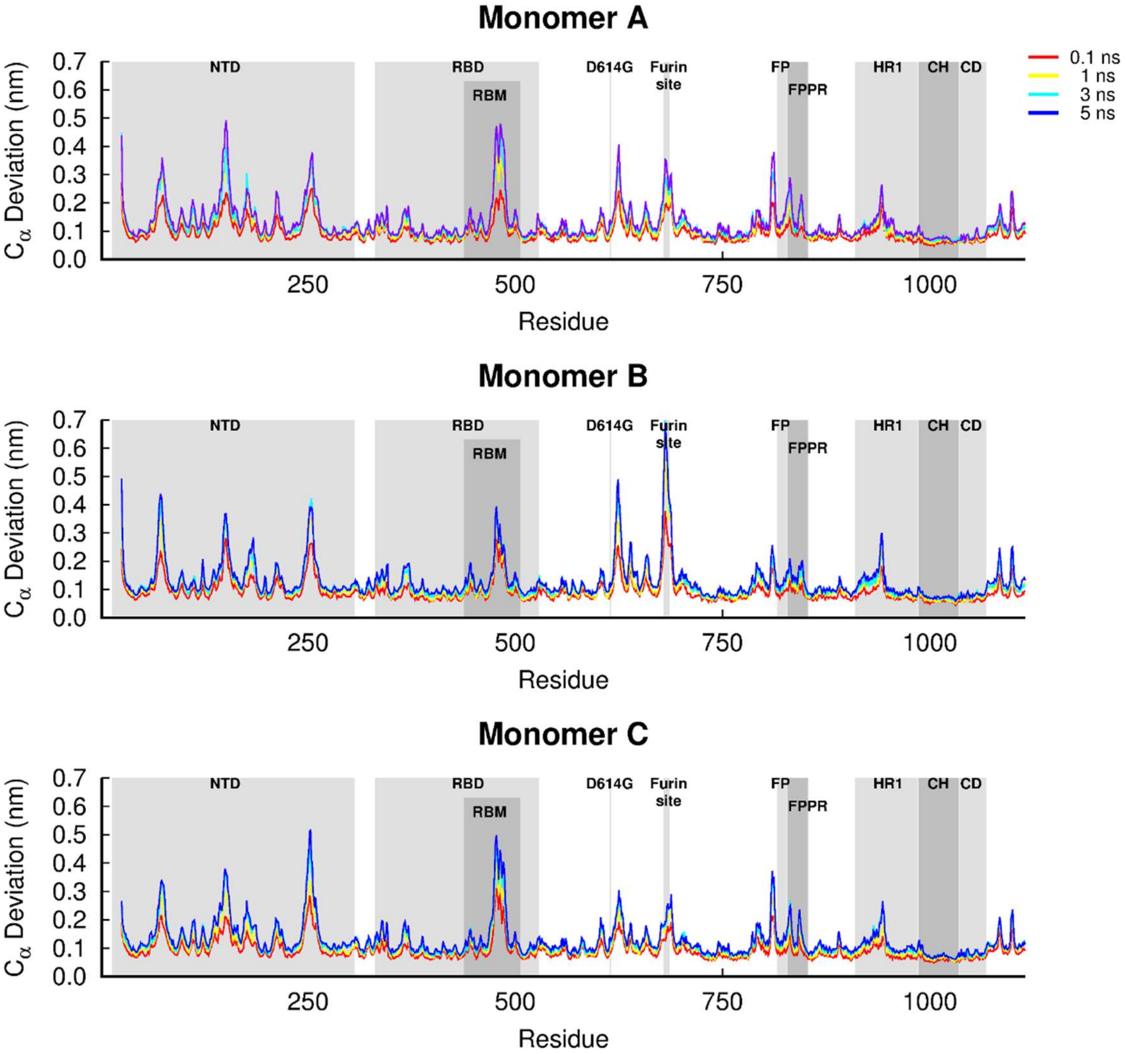


**Figure S19.** Average Cα-positional deviations in the five nanoseconds after LA removal from the FA sites in the D614G spike from SARS-CoV-2. The average deviations were calculated using the Kubo-Onsager approach (*13-15*) for the pairwise comparison between the short apo D614G simulations and the LA-bound D614G simulations. The positions of some important structural motifs are highlighted in grey, namely N-terminal domain (NTD), receptor-binding domain (RBD), receptor-binding motif (RBM), fusion peptide (FP), fusion-peptide proximal region (FPPR), heptad repeat 1 (HR1), central helix (CH), connector domain (CD). Please zoom in to the image for detailed visualisation

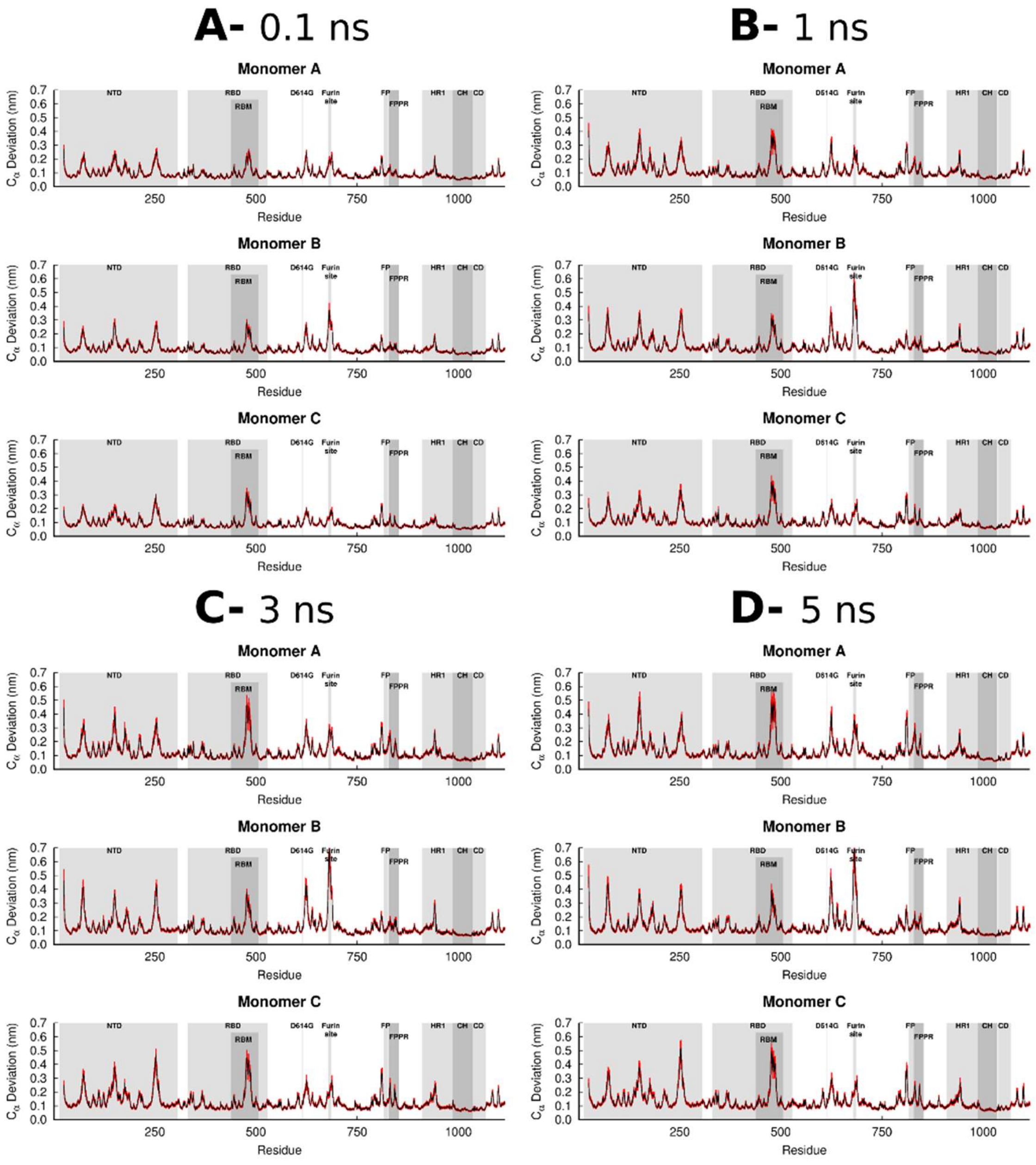


**Figure S20.** Average Cα-positional deviation (and corresponding standard errors) in the 0.1 (**A**), 1 (**B**), 3 (**C**) and 5 (**D**) ns after LA removal from the FA site in the D614G spike. The average deviations were calculated using the Kubo-Onsager approach (*13-15*) for the pairwise comparison between the nonequilibrium apo D614G simulations and equilibrium LA-bound D614G simulations. The vertical red lines represent the standard error of the mean. The positions of some important structural motifs are highlighted in grey, namely N-terminal domain (NTD), receptor-binding domain (RBD), receptor-binding motif (RBM), fusion peptide (FP), fusion-peptide proximal region (FPPR), heptad repeat 1 (HR1), central helix (CH), connector domain (CD). Please zoom in to the image for detailed visualisation

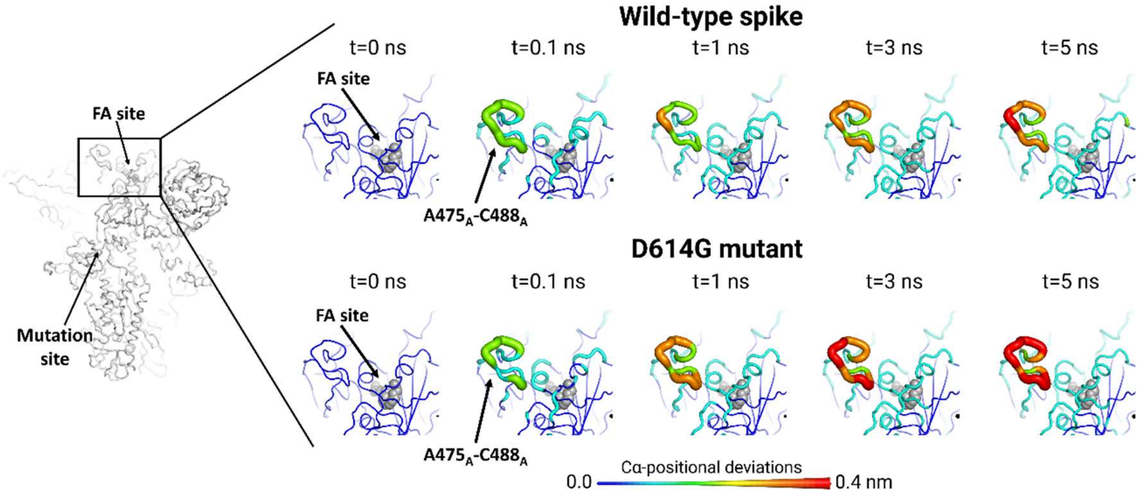


**Figure S21.** Differences in the conformational response of the $RBD_A$ to LA removal between the wild-type and D614G spike. Average Cα-positional deviation around the second FA site at times 0, 0.1, 1, 3 and 5 ns following LA removal from the FA binding pockets. The Cα deviations between the simulations with and without LA were determined for each residue, and the final values were averaged over the 90 pairs of simulations (Figures 2, S4 and S19). The Cα average deviations are mapped onto the structure used as the starting point for the LA-bound simulations. Note that in this image, both structure colours and cartoon thickness relates to the average Cα-positional deviation values. LA molecule is represented with grey spheres. The subscript letters in the labels correspond to the monomer ID.

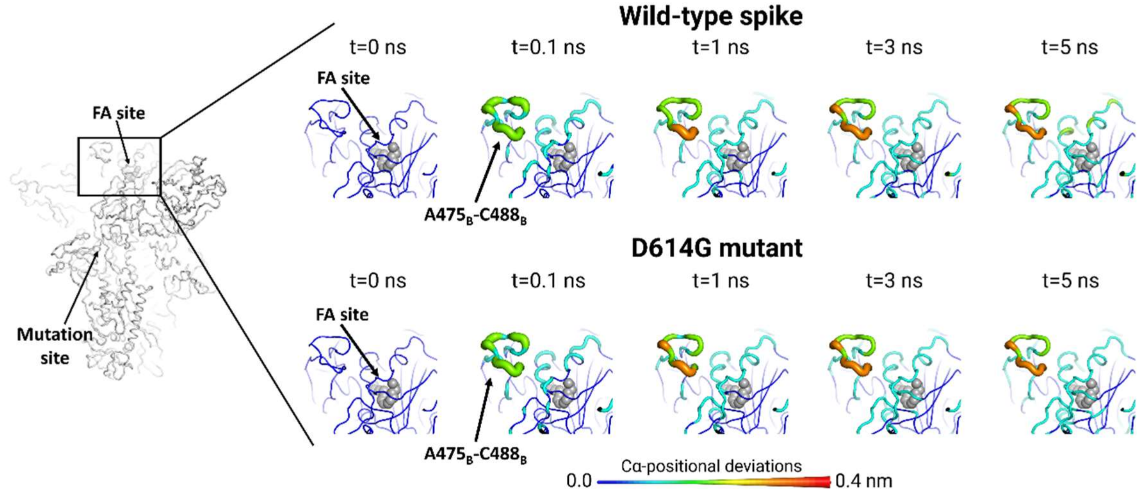


**Figure S22.** Differences in the conformational response of the $RBD_B$ to LA removal between the wild-type and D614G spike from SARS-CoV-2. Average Cα-positional deviation around the third FA site at times 0, 0.1, 1, 3 and 5 ns following LA removal from the FA binding pockets. For more details, see the legend of Figure S21.

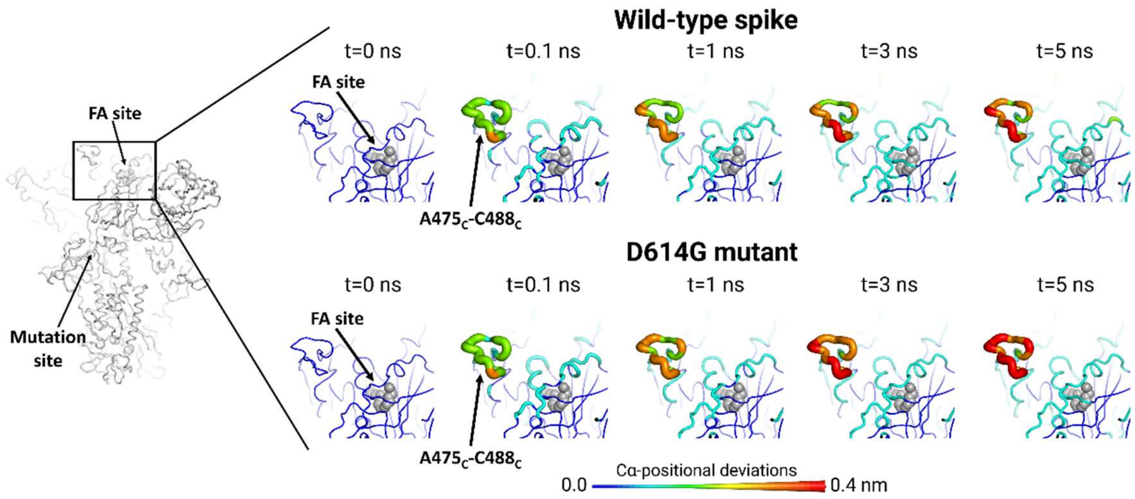


**Figure S23.** Differences in the conformational response of the $RBD_C$ to LA removal between the wild-type and D614G spike from SARS-CoV-2. Average Cα-positional deviation around the first FA site at times 0, 0.1, 1, 3 and 5 ns following LA removal from the FA binding pockets. For more details, see the legend of Figure S21.

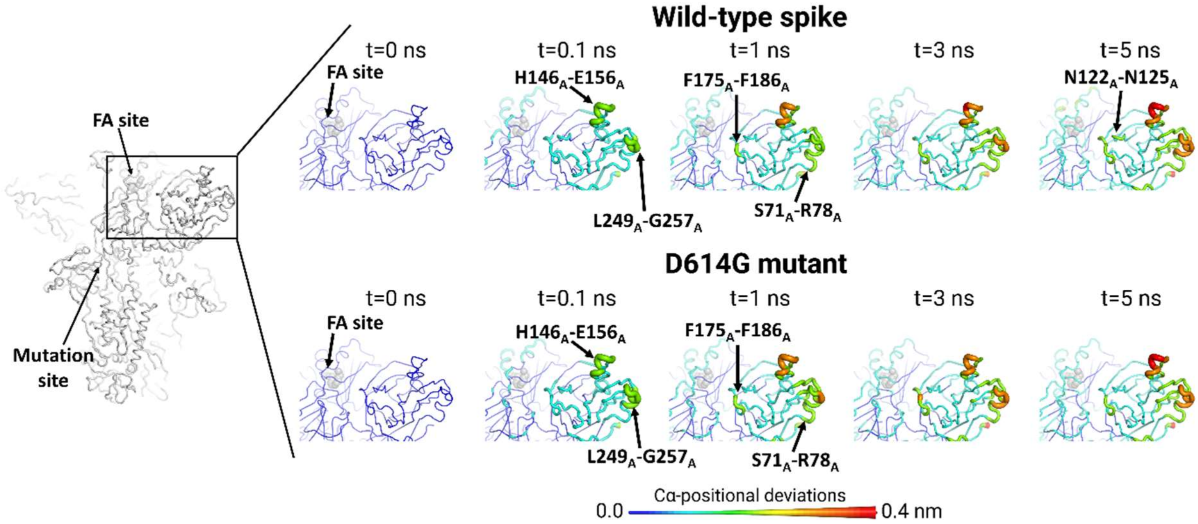


**Figure S24.** Differences in the conformational response of the $NTD_A$ to LA removal between the wild-type and D614G spike from SARS-CoV-2. Average Cα-positional deviation around the third FA site at times 0, 0.1, 1, 3 and 5 ns following LA removal from the FA binding pockets. For more details, see the legend of Figure S21.

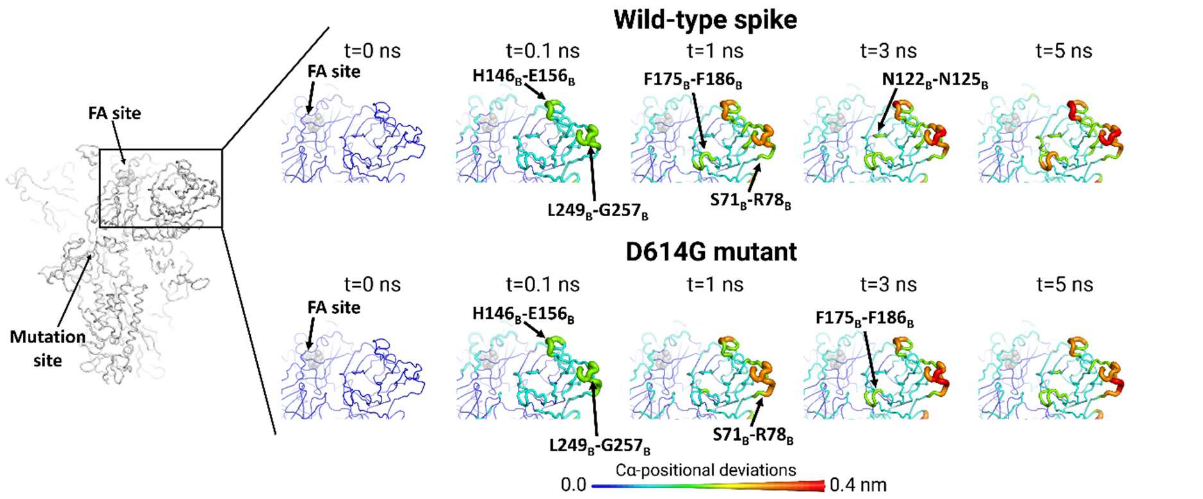


**Figure S25.** Differences in the conformational response of the NTD$_B$ to LA removal between the wild-type and D614G spike from SARS-CoV-2. Average Cα-positional deviation around the first FA site at times 0, 0.1, 1, 3 and 5 ns following LA removal from the FA binding pockets. For more details, see the legend of Figure S21.

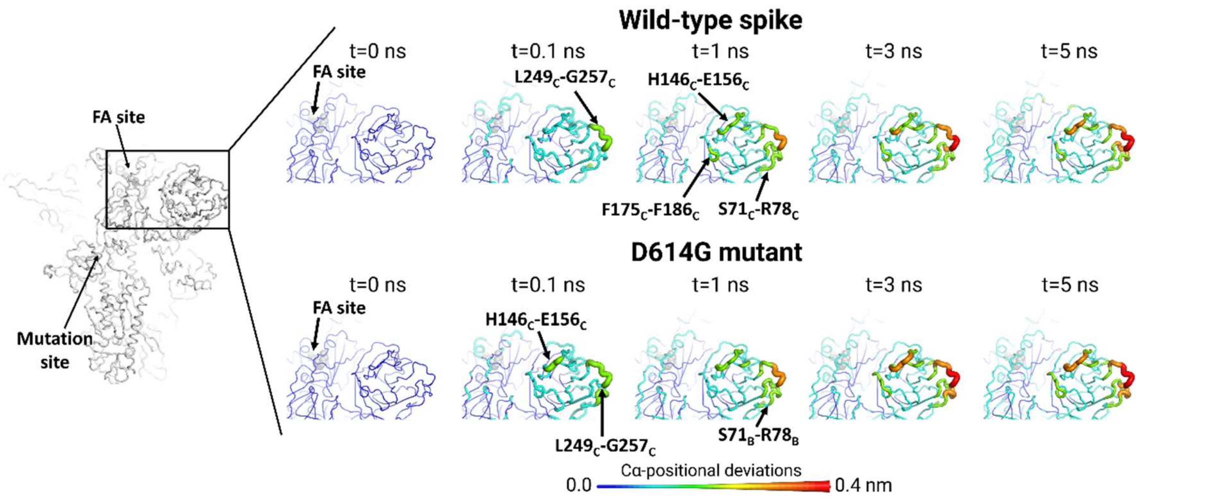


**Figure S26.** Differences in the conformational response of the NTD$_C$ to LA removal between the wild-type and D614G spike from SARS-CoV-2. Average Cα-positional deviation around the second FA site at times 0, 0.1, 1, 3 and 5 ns following LA removal from the FA binding pockets. For more details, see the legend of Figure S21.

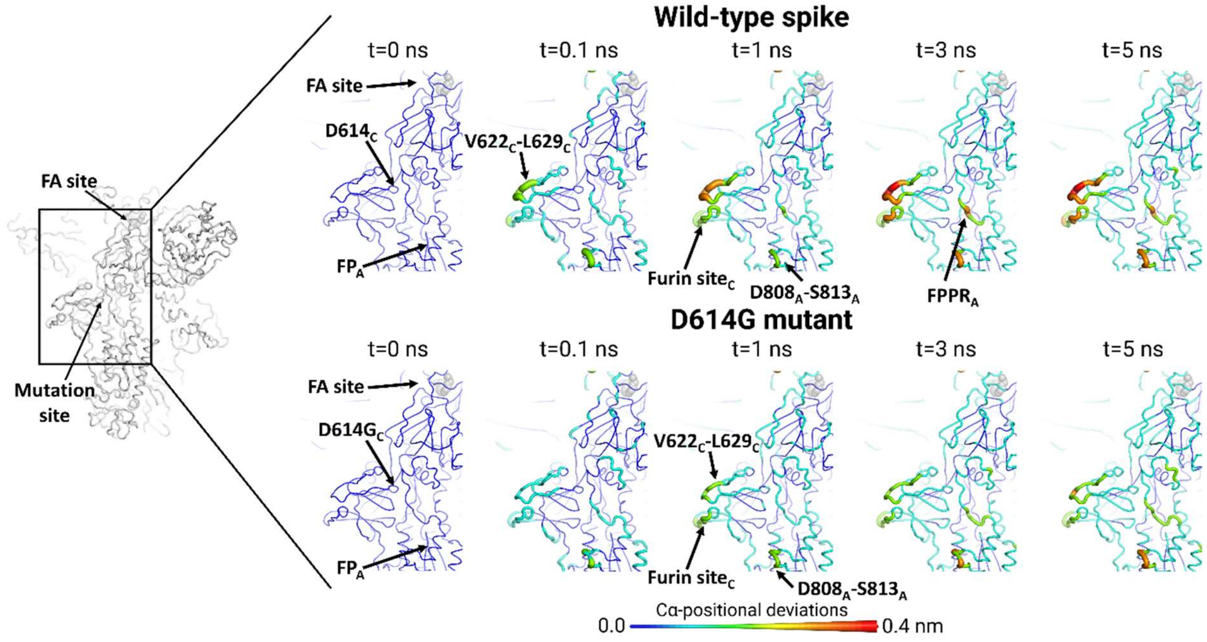


**Figure S27.** Differences in the conformational response of the $FPPR_A$ to LA removal between the wild-type and D614G spike from SARS-CoV-2. Average Cα-positional deviation around the third FA site at times 0, 0.1, 1, 3 and 5 ns following LA removal from the FA binding pockets. For more details, see the legend of Figure S21.

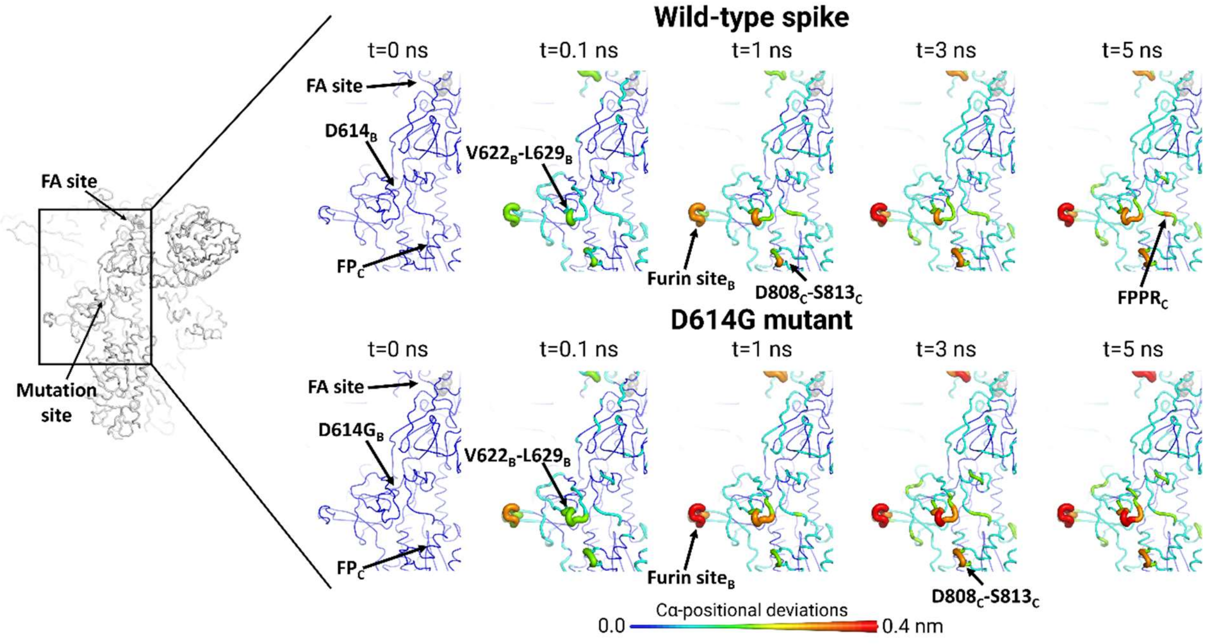


**Figure S28.** Differences in the conformational response of the $FPPR_C$ to LA removal between the wild-type and D614G spike from SARS-CoV-2. Average Cα-positional deviation around the second FA site at times 0, 0.1, 1, 3 and 5 ns following LA removal from the FA binding pockets. For more details, see the legend of Figure S21.

### Supporting movies

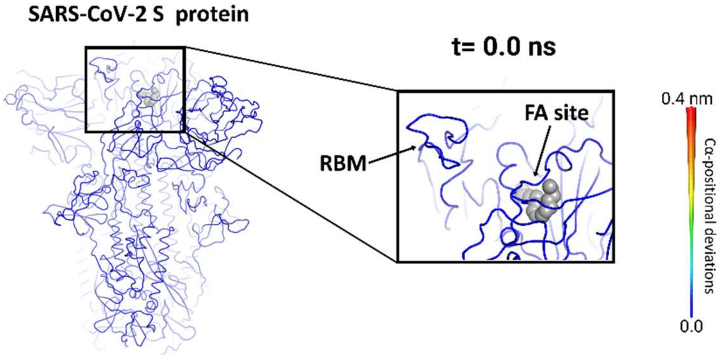


**Movie1.** Communication pathways between the FA site and RBD, NTD, furin and FP sites in the wild-type spike protein from SARS-CoV-2 (Signal_propagation_in_wildtype.mp4).

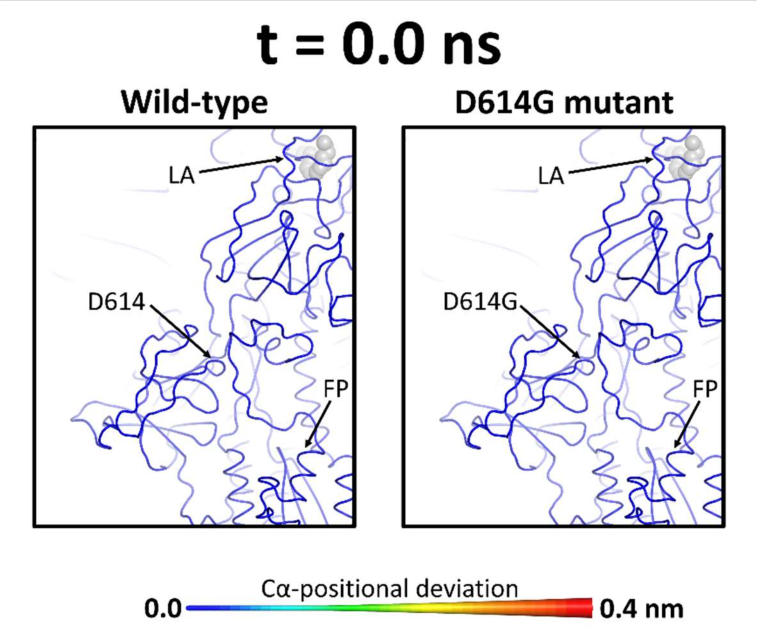


**Movie2.** Comparison between the communication pathways in the wild-type and D614G spike from SARS-CoV-2 (Comparison_wildtype-D614G.mp4).
